# Genetically encoded mechano-sensors with versatile readouts and compact size

**DOI:** 10.1101/2025.01.16.633409

**Authors:** Yuan Ren, Jie Yang, Takumi Saito, Oliver Glomb, Avinash Kumar, Sayed Iman Mousavi, Brigitte Naughton, Christina de Fontnouvelle, Barbara Fujita, Zhiqun Xi, Christian Schlieker, Shaul Yogev, Yongli Zhang, Julien Berro

## Abstract

Mechanical forces are critical for virtually all fundamental biological processes, yet quantification of mechanical forces at the molecular scale *in vivo* remains challenging. Here, we present a new strategy using calibrated coiled coils as genetically encoded, compact, tunable, and modular mechano-sensors to substantially simplify force measurement *in vivo*, via diverse readouts (luminescence, fluorescence and analytical biochemistry) and instrumentation readily available in biology labs. We demonstrate the broad applicability and ease-of-use of these coiled coil mechano-sensors by measuring forces during cytokinesis (formin Cdc12) and endocytosis (epsin Ent1) in yeast, force distributions in nematode axons (β-spectrin UNC-70), and forces transmitted to the nucleus (mini-nesprin-2G) and within focal adhesions (vinculin) in mammalian cells. We report discoveries in intracellular force transmission that have been elusive to existing tools.

## Main

Currently there are two major categories of sensors to measure mechanical forces at the single molecule level for live biological samples. One uses DNA duplexes (hairpins) that rupture at different levels of force^1–3^, and the other one utilizes elastic peptides (nano-springs) that monotonically change their extension in a force-dependent manner^4–6^ (Fig. 1a-b). Both are tuned to maximize their conformational change to tensile mechanical forces in the pico-newton (pN) range, and the conformational changes are commonly quantified by Förster Resonance Energy Transfer (FRET) measurements^7–10^. Variations of mechano-sensors based on these principles have had great successes and yielded valuable insights into the mechanical workings of cell attachment, cell motility, and embryo development^11–16^. Nevertheless, both approaches have inherent limitations in their broad applications. For the DNA duplexes, it is easy to tune their mechanical properties and alter readouts^17–21^ (Fig. 1a). However, it is difficult to deliver DNA duplexes into cells and protect them from rapid degradation. Therefore, in practice, DNA duplexes are only applied to measure force on proteins with extracellular domains^10,22^. Mechano-sensors based on calibrated elastic peptides, on the other hand, can be genetically encoded within any protein of interest, in theory. However, the inclusion of two fluorescent proteins for FRET measurement results in a large sensor size (>55 kDa) that is rarely tolerated by the host protein (Fig. 1b). Additionally, FRET measurements are inherently challenging to perform, requiring extensive controls and calibration^23,24^. These measurements are highly context-dependent, time-consuming, and exhibit a low to moderate dynamic range (5-25%) *in vivo*^23–26^. Signals from FRET-based tension sensors are typically averaged over a population of molecules from unknown distribution and quickly drops to a flat value outside of the sensors’ linear range, resulting in convolution of the forces on single molecules. Although newer versions of FRET-based sensors have improved performance by introducing digital force responses and multiplexed pairs^12,13^, smaller genetically encoded mechano-sensors that do not rely on FRET measurements are still desired to democratize molecular force measurements in cell and developmental mechanobiology beyond the few commonly studied molecules.

**Figure 1.**
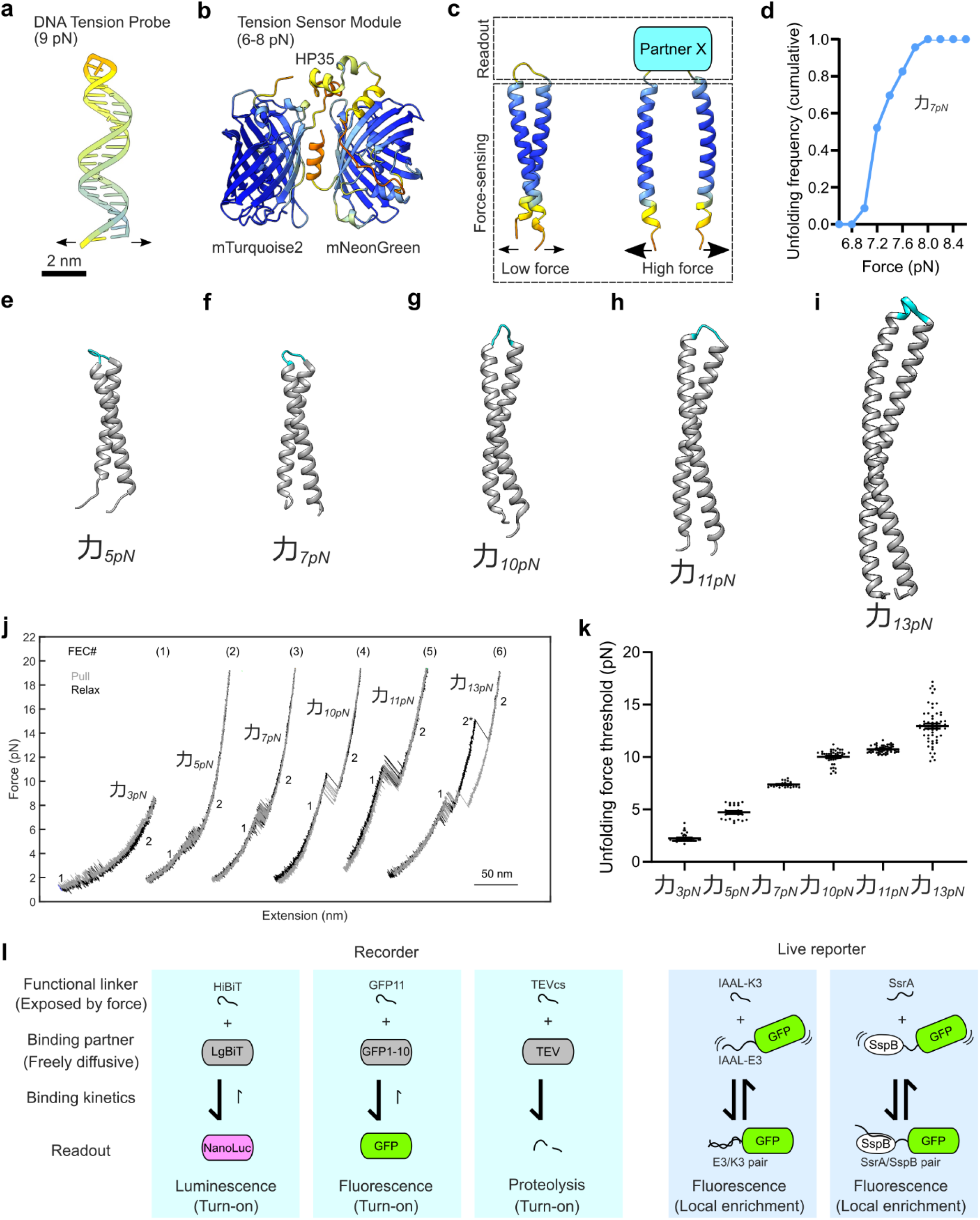
Rational design and calibration of coiled coils as force-sensing modules. **a**, A DNA hairpin (TP9) with defined nucleotide composition unfolds at a characteristic force threshold^2^. Fluorescence readouts (such as a fluorophore-quencher pair) can be added to nucleotides by chemical modifications on the DNA hairpin so that change in fluorescence reports the digital opening of the DNA under force. **b**, An elastic peptide (HP35) sandwiched by two fluorescent proteins (mTurquoise2 and mNeonGreen) form a genetically encoded force sensor^16,30^. The FRET efficiency between the two fluorescent proteins depends on the peptide length, which scales linearly with force magnitude within a small range (e.g. 3-6 pN). **c**, Calibrated coiled coils can be used as force-sensing modules to correlate mechanical force to their conformational change. Readouts are protein constructs that can only bind to the linkers connecting the two α-helices of the coiled-coil when the coiled-coil is unfolded under force. **a**-**c**, Molecules are shown at the same scale, and the structures are predicted by AlphaFold3^103^ and colored according to the residues’ b-factors. Arrows indicate the directions of force pulling. **d**, Cumulative unfolding probability of the coiled coil force sensor 力_7*pN*_ as a function of force. The jump of the unfolding probability from below 0.1 to above 0.9 within 1 pN indicates a near-digitalresponse to mechanical forces. **e-i**, Predicted structures of 力_5*pN*_, 力_7*pN*_, 力_10*pN*_, 力_11*pN*_ and 力_13*pN*_with their GS linkers (GGSSGG) highlighted in cyan are shown at the same scale as **a**-**c**. Mechanical stabilities are tuned by mutating the hydrophobic core (amino acids in positions *a* and *d*) and by changing the total number of heptads. See also Fig. S1 for helical wheel depictions. **j**, Representative force-extension curves (FECs) obtained by pulling (grey) or relaxing (black) coiled coil sensors using optical tweezers. The conformations of the coiled coils are labeled as follows: 1, folded state; 2, unfolded state; and 2*, partially unfolded state. **k**, Unfolding force thresholds of coiled coils are calibrated by optical traps. Each dot represents a pulling event. **l**, The selection of functional linkers and their binding partners offers modularity in force sensor readouts. Readouts with slow unbinding kinetics are force recorders (HiBiT and LgBiT; GFP11 and GFP1-10; TEV cleaving site (TEVcs) and TEVp), while readouts with fast unbinding kinetics are force live reporters (IAAL-K3 and IAAL-E3; SsrA and SspB).

We present a new class of mechano-sensors that retains the programmable digital mechanical response and versatile readouts of DNA sensors, while being fully genetically encoded and substantially smaller than the current peptide-based force sensors^24,26^ (Fig. 1c). We achieve this goal through a modular approach by building on force sensors we have previously developed^27–29^ (Fig. S11). The *force-sensing module* is composed of a dimeric anti-parallel coiled coil that unfolds (i.e. fully opens) when the tensile force applied to it is larger than a calibrated force threshold. The *readout module* is bipartite and made of a) a peptide that links the two α-helices of the coiled coil and b) an interacting partner that binds the linker only when the coiled coil is in an open conformation but not when it is in a closed conformation (Fig. 1c). The binding between different linkers and their binding partners can be engineered to generate binary signals that are gated by the force on the coiled coil (Fig. 1l). Due to their small size and short end-to-end distances (∼13 Å, or the typical length of a ∼5 amino acid linker), the mechano-sensors can be placed between virtually any domains or within virtually any flexible region of a protein of interest with minimal influence on the protein’s function. Our approach extends and generalizes previous analogous approaches based on larger force-sensing modules, including mutants of the HP35 peptide or Titin I10 domain, which have been limited to proteins in focal adhesions (talin and vinculin)^30–32^.

We derived our library of force-sensing modules from an artificial heterodimeric antiparallel coiled coil^33^, which unfolds at 7.4 ± 0.1 pN (mean ± SEM) (Fig. 1d). Dimeric coiled coils are stabilized by knobs-into-holes packing at the hydrophobic core along with electrostatic pairing on the surface^34,35^. We tuned the mechanical stability by 1) varying the amino acids in the hydrophobic core and 2) changing the total number of heptad repeats (Fig. 1e-I, S1a-e). We also used molecular simulations to aid our design (Fig. S1f). The resultant coiled coils have the same end-to-end distance and surface properties when folded, and each sensor reversibly unfolds when the tensile force exerted on it exceeds a predetermined sequence-specific threshold between 5 pN (four heptads) and 13 pN (eight heptads) (Fig. 1j-k). As shown in Fig. 1j, the folded coiled coils start to switch (or flicker at equilibrium) between folded and unfolded conformations when force reaches the threshold (Fig. 1e, state 1 and state 2, respectively), and become completely unfolded when force keeps increasing. The unfolding/refolding transitions are reversible in the relaxation rounds (Fig. 1j, black curves). We took this reversible transition feature as a main criterion to select sensors for further use in live measurements. The lower bound of the unfolding force threshold is set by the minimum number of heptads required to form a coiled coil (3.5 heptads for most known anti-parallel coiled coils)^36,37^. Higher force thresholds can be achieved by increasing the number of heptads but with diminishing returns the longer it is extended. The maximum number of heptads is also limited by the increasing likelihood of unfolding hysteresis when the number of heptads exceeds 6 (Fig. 1j, FEC# (6)), suggested by the additional folding intermediate (Fig.1j, state 2*). To detect forces smaller than 5 pN, we engineered the stalk region of mouse cytoplasmic dynein, an anti-parallel coiled coil that unfolds at 3 pN (Fig. 1j, FEC # (1)). Our library of four force-sensing coiled coils (hereafter called 力_*XpN*_, where 力 (lì) is the Chinese character for “force” and X pN is the opening force threshold) is sufficient to cover the physiological range of most forces on a single molecule with ∼2 pN resolution. Note that varying the pulling speed or the replacement of the linkers of similar lengths between the two α-helices has a minimal influence on the unfolding force threshold of the coiled coil, consistent with previous models and measurements^38,39^ (Fig. S2).

We developed two categories of readout modules: *recorders* and *live reporters*. “Recorders” use bipartite systems that are irreversible (or very slowly reversible), so that they keep emitting signals for a long time after the force-sensing module has unfolded, and therefore “records” (or “integrates”) the maximum force the sensor has experienced in the past (Fig. 1l). Here, we present three recorders: the split-Nanoluc, split-GFP, and TEV systems. The split-Nanoluc system is composed of an 11-aa (amino acid) peptide (HiBit) used as linker between the two α-helices, and the rest of the Nanoluc (LgBit) expressed in the cell^40,41^. When force opens the force-sensing element, the exposed HiBit is bound by LgBit, reconstituting a full Nanoluc able to produce luminescence following the addition of a Nanoluc substrate (e.g. furimazine). Similarly, the split-GFP system is composed of GFP11, a 16-aa peptide whose sequence corresponds to the 11^th^ β-strand of the GFP, used as a linker between the α-helices of the force-sensing module, and GFP1-10, the GFP lacking the 11^th^ β-strand (not fluorescent by itself), which is expressed in the cell. When force is low, no fluorescence is emitted. Upon opening of the force-sensing element, GFP11 is exposed and binds with GFP1-10, matures, and emits fluorescence^42^. Because the off-rate constant of the split-GFP system is very slow (∼hours), fluorescence persists even after the force vanishes^43^. Since sub-cellular compartments may limit the use of fluorescence or luminescence, we developed an orthogonal detection method based on cleavage by Tobacco Etch Virus (TEV) protease (TEVp). The TEV readout uses as a linker the 7-aa TEV cleavage sequence (TEVcs) which can be cleaved by the TEVp. In a cell expressing the TEVp, the force sensing element is cleaved after it is unfolded by force^44^, and the cleavage can typically be detected using an immunoblot.

“Live reporters” are constituted of a peptide tag used as linker and a complementary peptide or domain fused to a fluorescent protein, which is expressed in the cell (Fig. 1l). Here, we present the reversible binder-tag systems IAAL-E3/IAAL-K3 and SsrA/SspB, an artificial binding pair^45^. Under low force, fluorescence is diffuse, but when high force unfolds the force-sensing element, the fluorescent protein re-localizes to the force sensor. Because the tags used are reversible with a fast off rate constant (see *in vivo* binding kinetics in Fig. 3j), when force is released, the fluorescent protein detaches, and fluorescence becomes diffuse again^46^. We summarized the strengths and weaknesses of each readout in Table 1, and simulated the kinetics of reporter binding in Note S1. Hereafter, the force-sensing coiled coils are called 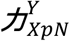, where X is the force threshold of the sensor and Y is the linker specific to the readout used to detect its opening (e.g. 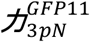 is the force sensor that opens under forces larger than 3 pN with the split-GFP readout)

**Table 1.**
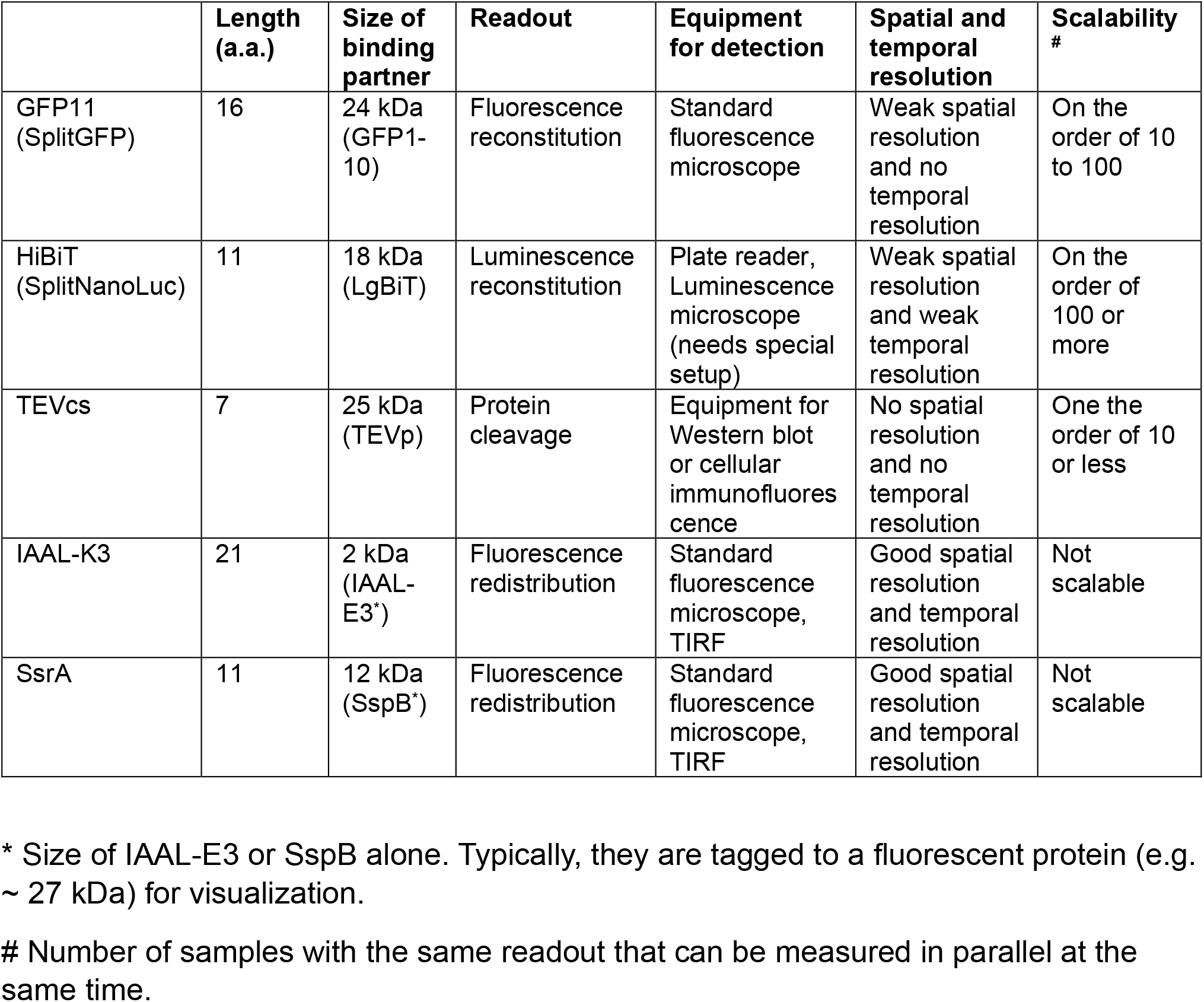
Comparison of functional linkers for force measurement.

We demonstrate the use of our new sensors by measuring the *in vivo* forces on key force-bearing cytoplasmic proteins. First, we measured the force on Cdc12, the formin that polymerizes actin filaments and connects them to the plasma membrane during cytokinesis in the fission yeast *Schizosaccharomyces pombe*^47,48^. Using the split-Nanoluc (Fig. 2b, Fig. S3) and split-GFP (Fig. 2c, Fig. S4) recorder readouts, we measured 6 pN peak tension on formin Cdc12 (Fig. 2a). Next, we followed the temporal evolution of the force on Cdc12 over the course of cytokinesis using the live reporter readout IAAL-K3/ mEGFP-IAAL-E3 (Fig. 2d). Timelapse imaging indicates that force on Cdc12 starts to exceed 3 pN during ring constriction and drops below 3 pN before the ring fully disassembles, suggesting that the mechanical tension on Cdc12 is down-regulated to match the decreased need for F-actin assembly. These results are the first force measurements on a formin *in vivo*. Previous *in vitro* measurements using a microfluidic chamber showed that weak tension on budding yeast formin Bni1 or mammalian formin mDia1 accelerates the formin-mediated polymerization of profilin-actin^49,50^. In addition, *in vitro* single-molecule spectroscopy experiments demonstrated that ∼7 pN tension on formin mDia1 accelerates its actin-polymerizing activity by nearly 8-fold, in the presence or absence of profilin^51,52^. Our *in vivo* force measurement of 6 pN on native Cdc12 strongly suggests that mechanical force regulates the activity of formin during cytokinesis^53^.

**Figure 2.**
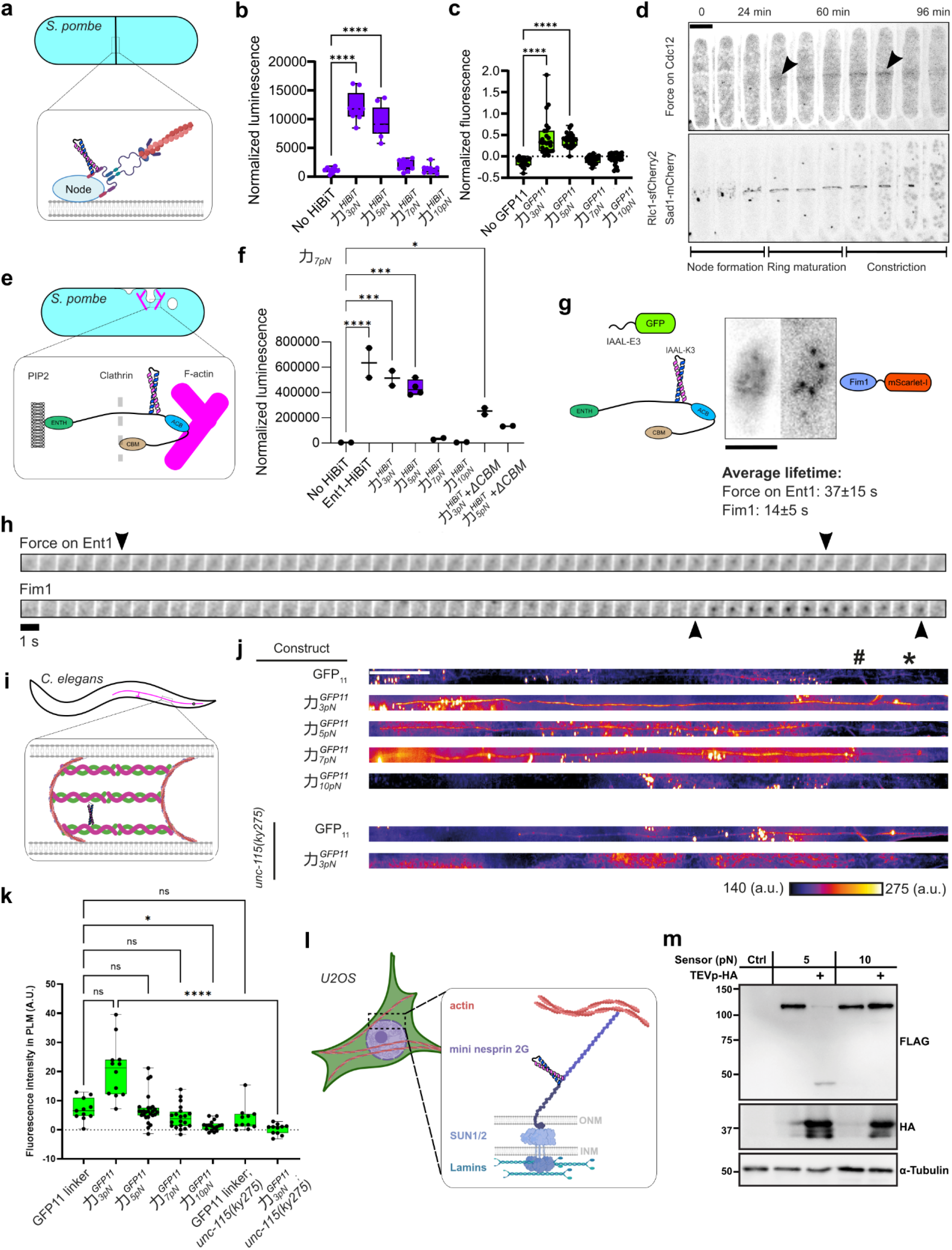
Modular assembly of mechano-sensors with versatile readouts in multiple biological systems. **a**, Schematic of the role of Cdc12 during cytokinesis in the fission yeast *S. pombe*. Cdc12 nucleates and polymerizes actin filaments, and bridges F-actin to the cytokinetic nodes on the plasma membrane. Force sensors are inserted after A216 of cdc12 genomic location. **b**, Force measurement on Cdc12 using the split-NanoLuc readout. Ordinary one-way ANOVA was performed with Tukey’s multiple comparison tests with “No-HiBiT” as the control and only displayed for pairs where p values is less than 0.05. ****, p<0.0001. Each dot represents one measurement with >1000 cells. Data are pooled from three independent repeats. **c**, Force measurement on Cdc12 using the split-GFP readout. Ordinary one-way ANOVA was performed with Tukey’s multiple comparison tests with “No-GFP11” as the control and only displayed for pairs where p values is smaller than 0.05. ****, p<0.0001. Each dot represents measurement from a single cell. Data are pooled from more than three independent experiments. **d**, Timelapse of force on Cdc12 (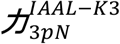 and mEGFP-IAAL-E3 binder) and the cytokinetic ring (Rlc1-sfCherry) during fission yeast cytokinesis. Sad1-mCherry is used to locate the dividing spindle pole body and to time cytokinesis. Force on Cdc12 starts to build up above 3 pN at the beginning of ring maturation and drops below 3 pN before the ring fully disassembles. Arrowheads indicate the recruitment of mEGFP to Cdc12 when 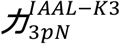 is unfolded by force. Scale bar, 5 μm. **e**, Schematic of Ent1 and its interacting partners during clathrin-mediated endocytosis (CME) in the fission yeast. Force sensors are inserted after P571 at ent1 genomic location. ENTH, N-terminal lipid-binding domain. ACB, actin cytoskeleton-binding domain. CBM, clathrin-binding motif. **f**, Force measurement on Ent1 using the split-NanoLuc readout. ∼6 pN force is detected on Ent1 and the deletion of CMB decreased the luminescence signal detected, indicating a smaller fraction of Ent1 molecules under force. Ordinary one-way ANOVA was performed with “No-HiBiT” as the control and only displayed for pairs where p values is smaller than 0.05. *, p<0.05; **, p<0.01; ***, p<0.001; ****, p<0.0001. Each dot represents one independent experiment with >1000 cells. See also Fig. S5 for controls and force measurement on different sites of Ent1. **g**, Dual-color TIRF is used to simultaneously track the force on Ent1 (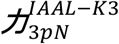 and mEGFP-IAAL-E3 binder) and actin dynamics (Fim1-mScarlet-I) in fission yeast cells. Scale bar, 5 μm. **h**, Montage of a representative CME event. Note that force on Ent1 increases to above 3 pN ∼30 s before actin assembly begins. Arrowheads denote the first and last frames where force on Ent1 is above 3 pN. **i**, Position of the sensory PLM neuron in the nematode *C. elegans*. Spectrins (UNC-70 and SPC-1), shown as spirals in the zoomed region, form the central building block of the membrane associated periodic skeleton (MPS), which consists of actin rings that are interspaced by spectrin tetramers below the plasma membrane throughout the entire length of the axon. Force sensors were inserted into the genomic unc-70 locus between spectrin repeats 8 and 9 (after R1167). Only one coiled coil is shown for clarity. See Fig. S6 for details of force sensor insertion. **j**, Representative maximum projections of *C. elegans* strains expressing UNC-70 with force sensors together with GFP1-10 under a PLM specific promoter (mec-17p) in wildtype or in a loss of function mutant of unc-115(ky275). Signal intensity is color coded according to the displayed color bar. Scale bar, 50 μm. The position of the cell body (*) and the rectum (#) are indicated. Background fluorescence outside of the axon region was not included in quantification. Example images are rescaled in X and Y dimension (1:2) to enlarge the neurite diameter for better visualization. **k**, Quantification of fluorescence in **j**. Multiple comparisons with the Kruskal-Wallis test. *, p<0.05. ****, p<0.0001. Each dot represents measurement from a single axon. **l**, Mini-Nesprin-2G transmits forces from cytoplasmic actin filaments to SUN in the nucleus. 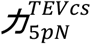 and 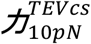 were inserted between the N-terminal filamentous actin-binding CH domain (1-485) and the C-terminal KASH SUN-binding domain (6525–6874). **m**, Immunoblot of cell lysates from U2OS cells transfected with 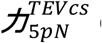 or 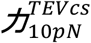 in mini-Nesprin-2G. Note that cleavage of 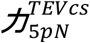 is observed upon co-transfection with TEVp, whereas 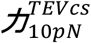 is resistant to TEVp activity (TEVp-HA +). Representative Western blot of two independent repeats. All schematics in this figure are for illustrative purpose only and not drawn to scale. Some schematics were created with BioRender.com.

Second, we measured the forces on epsin Ent1, an adaptor protein involved in clathrin-mediated endocytosis. Ent1 contains an actin cytoskeleton-binding (ACB) domain and a clathrin-binding motif (CBM)^54,55^ (Fig. 2e). Although the ACB domain has been shown to be critical for connecting the plasma membrane to the actin cytoskeleton^55,56^, we suspected that the CBM may also contribute to the force relay due to the positioning of the clathrin coat within the F-actin layer (Fig. 2e). Indeed, using the split-Nanoluc readout, we detected a peak force of 6 pN before the ACB domain (at P571) (Fig. 2f). Importantly, elimination of the C-terminal CBM decreased the force by 75%, indicating that substantial tension on Ent1 is mediated by its interaction directly with the clathrin lattice, and not just from the actin cytoskeleton, as previously thought^27,55^ (Fig. 2f, Fig. S5). We corroborated this conclusion by detecting forces above 3 pN both before and after the ACB domain (at A637), using the split-GFP readout as well as the TEVp readout (Fig. S5d-k). In agreement with this result, dual-color TIRF imaging of the force on Ent1 and the F-actin crosslinking protein fimbrin, which is used as a fiduciary marker of filamentous actin at endocytic sites (Fig. 2g), indicates that force on Ent1 starts to accumulate ∼30 seconds before F-actin starts assembling^57,58^ (Fig. 2h, Fig. S5l-n). This surprising result directly demonstrates for the first time that forces during endocytosis are not exclusively produced by actin assembly^55,59,60^. Actin may interact with Ent1 at the later stages of endocytosis to pull the Ent1 tail from the clathrin layer to the actin layer^61^.

Third, we inserted the force sensors using the split-GFP recorder readout into *C. elegans* β-spectrin (UNC-70), which forms rungs between periodic F-actin rings in the submembrane region of axons (termed membrane-associated periodic skeleton or MPS)^62,63^ (Fig. 2i, Fig. S6). We measured a peak force of ∼8 pN on β-spectrin, which was dependent on the integrity of the MPS lattice, as impairment of MPS lattice formation in *unc-115(ky275)* mutants was required to maintain tension^64–66^ (Fig. 2j-k). This result suggests that the maximal force experienced by a single β-spectrin in freely-moving worms is higher than that measured in paralyzed worms, where FRET-based force sensors reported an average force of 1.5 pN at the same location on β-spectrin^67^.

Fourth, we measured the force in the linker of nucleoskeleton and cytoskeleton (LINC) complex responsible for force transmission across the nuclear envelope and nuclear positioning^68–70^. To this end, we inserted the mechano-sensor into mini-Nesprin-2G, a smaller functional version of the Nesprin-2G protein that associates with actin on the cytosolic side and the SUN protein within the lumen of the nuclear envelope^71,72^ (Fig 2l). Upon co-transfection with the TEV protease, mini-Nesprin-2G with cc-5 pN was cleaved, while the mini-Nesprin-2G with cc-10 pN remained resistant to proteolytic cleavage (Fig. 2m, S7). We confirmed these measurements using the split-GFP and the split-Nanoluc readouts (Fig. S8). Our results are consistent with previous studies showing that mini-Nesprin-2G is under actin-dependent tension in the single pN range^73,74^.

To showcase the transformative power of our new sensor design, we measured the forces on one of the well-studied mechano-sensitive protein, vinculin, using multiple readouts to investigate its behavior in different settings. Vinculin is a key focal adhesion protein, which is believed to be under ∼2 pN average tension, as determined from previous ensemble measurements with FRET-based force sensors in adherent cells^4,11,75^ (Fig. 3a). However, vinculin is also known to be a mechanosensitive protein that changes conformation under 5-15pN force to modulate its activity (e.g. vinculin becomes an actin filament catch bond when under ∼8pN force)^76,77^. In addition, the average force reported by FRET-based sensors is likely an underestimate because any vinculin under forces far outside of the linear range of the FRET-sensor emits the same signal as vinculin under force at the sensors’ bounds^10,16,24,27^. Last, ensemble *in vivo* measurements using FRET sensors are unable to represent the distribution of forces on the vinculin (i.e. are all vinculin molecules under the same force? Or are there a few vinculin molecules under very large force while most are under very low force? Etc.). Using our sensors with the split-GFP recorder readout, we showed that the majority of vinculin molecules (∼90%) under tension are experiencing force above 5 pN, and that ∼10% bear forces above 10 pN (Fig. 3b-c, S9). Similar ratios were measured using the split-Nanoluc readout using newly adherent cells on different substrates (Fig. 3d-f). These results demonstrate that a small percentage of vinculin molecules mediate most of the tension between talin and F-actin^12,13^. Our data also demonstrate that fewer vinculin molecules were under tension on Matrigel than on glass, with a reduced fraction of vinculin molecules under tension above 5 pN (Fig. 3e-f). To test if substrate stiffness alone changes the force on vinculin, we plated cells with the split-Nanoluc sensors on a 96-well plate with varying stiffness in each well (0.1 kPa to 100 kPa) (Fig. 3g), which, to our knowledge, is the first high-throughput single molecule force measurement on vinculin. We did not detect a strong dependence of force on vinculin on substrate stiffness (Fig. 3h-i), in agreement with a previous force measurement with a smaller sample size^78^. From our measurements, the large fraction of vinculin molecules with force above 5 pN (∼70%) across stiffness is consistent with the *in vitro* activation force threshold of single vinculin molecules, a force level needed for the conformational change of vinculin to relieve self-inhibition and to bind F-actin^11,76,79,80^. The much smaller fraction of vinculin molecules with force above 7 pN (∼10%) implies that activated vinculin may buffer force, possibly mediated by the flexible linker between its head and tail domains or by disengaging from other force transmitting proteins^81,82^. To demonstrate that the binding and unbinding kinetics of our IAAL-E3/IAAL-K3 reporter system is fast (sub-second to second time scales), we used Fluorescence Recovery After Photobleaching (FRAP) to measure a recovery rate with a half-time of 1.6 s (Fig. 3j-k). We also demonstrated that the size of the fluorescent tags is not limiting the detection ability of our sensors in the crowded environment of focal adhesions (Fig. S10).

**Figure 3.**
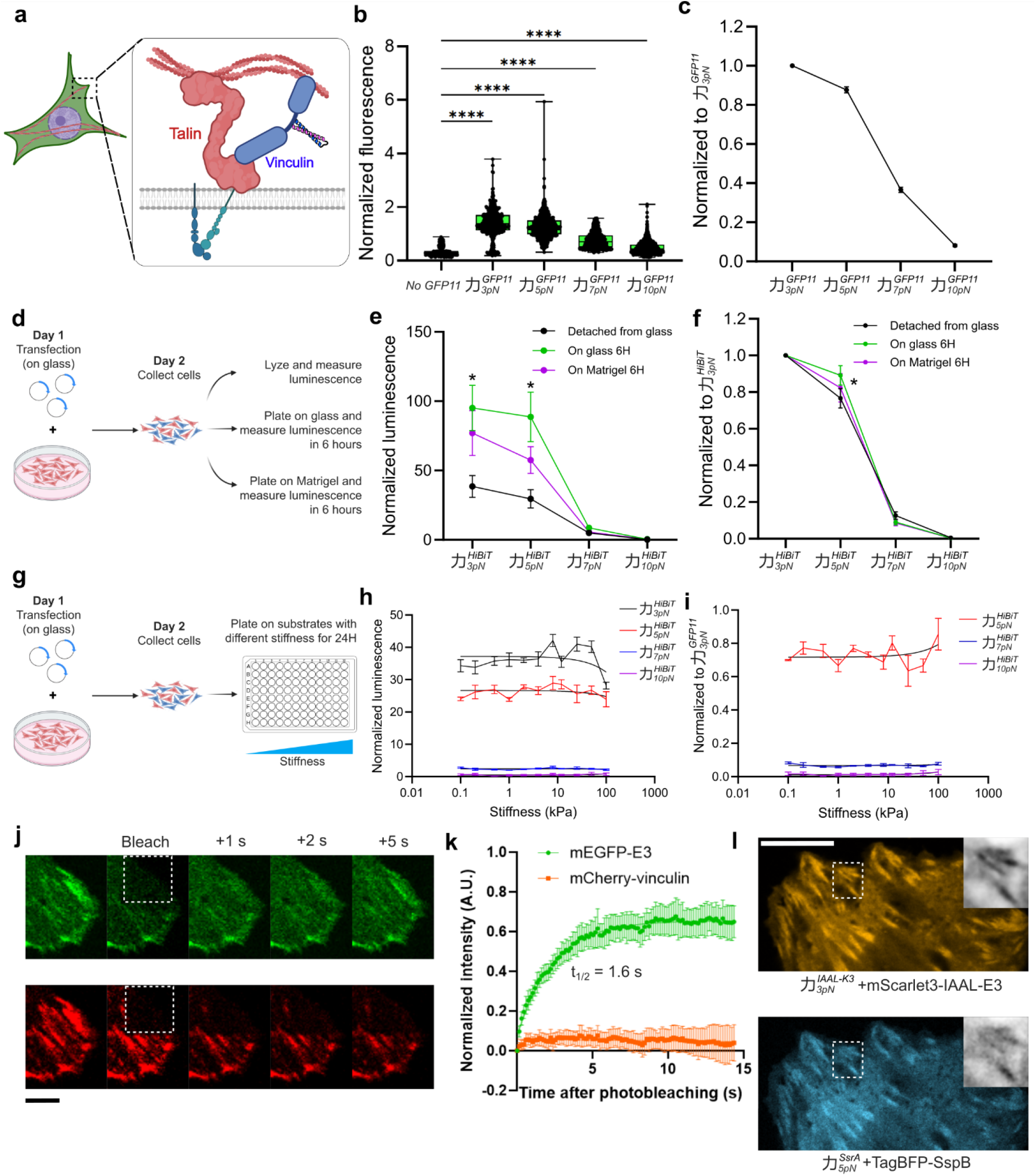
Comprehensive force measurements on vinculin. **a**, Vinculin binds to talin and F-actin in focal adhesions and contributes to force transmission between the extracellular matrix and the cytoskeleton. **b**, Quantification of force on vinculin with the Split-GFP readout. U2OS cells were plated on glass and transfected with plasmids encoding mCherry-vinculin with different force sensors inserted after E883 using the split-GFP readout. GFP fluorescence is normalized to mCherry-vinculin signals. Each dot represents measurement from a segmented focal adhesion. Data are pooled from two independent experiments. Ordinary one-way ANOVA was performed with Tukey’s multiple comparison tests with “No-GFP11” as the control. ****, p<0.0001. **c**, Force distribution of vinculin, calculated by normalizing the fluorescence signals from 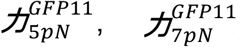 *and* 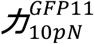 to that from 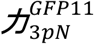. Mean ± SEM. **d**, Schematic for comparing the forces on vinculin from cells plated on glass or Matrigel for 6 hours. **e**, Quantification of force on vinculin with the Split-Nanoluc readout. Luminescence is normalized to mCherry-vinculin signals to compare the number of mechanically active vinculin molecules. Multiple comparisons with Kruskal-Wallis test. *, p<0.05. **f**, Force distribution on vinculin, calculated by normalizing the luminescence signals from different sensors to that from 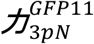. Multiple comparisons with Kruskal-Wallis test. *, p<0.05. **g**, U2OS cells were transfected with force sensors, collected, and plated onto substrates with different stiffnesses. Luminescence was measured to detect the forces on vinculin after overnight culture. **h**, Quantification of force on vinculin across substrate stiffness. Data are shown as mean ± SEM with fitted lines of linear regression. The slopes of the fitted lines are not significantly different from zero (F-test). **i**, Force distribution on vinculin calculated by normalization to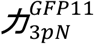. Data are shown as mean ± SEM with fitted lines of linear regression. The fraction of forces on vinculin was not changed by the substrate stiffness according to the F-test on the slopes of the fitted lines. **j**, U2OS cells were transfected by plasmids encoding mCherry-vinculin-E883-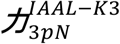 and mEGFP-IAAL-E3. FRAP was performed to compare the recovery of mEGFP and mCherry signals in focal adhesions. Scale bar: 5 μm. **k**, Quantification of fluorescence intensities after photobleaching. mEGFP signals recover with t_1/2_=1.6 s, indicating fast binding and unbinding kinetics for the live force reporter. **l**, U2OS cells were transfected with vinculin-E883-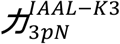 with mScarlet-3-IAAL-E3 binder, and vinculin-E883-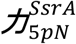 with TagBFP-SspB binder. Signals corresponding to 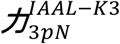 are closer to the cell edge and stronger in focal adhesions than 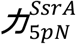. Scale bar, 10 μm. All schematics in this figure are for illustrative purpose only and not drawn to scale. Some schematics were created with BioRender.com.

To gain a spatial understanding of the forces on vinculin, we used two pairs of orthogonal binder/tag (IAAL-E3/IAAL-K3 and SsrA/SspB) to label mechanically active vinculin molecules and detect two levels of force (respectively 3 pN and 5 pN) with two different colors (respectively mScarlet-I and TagBFP) (Fig. 3l). Consistent with the split-GFP and split-Nanoluc readouts, we measured a smaller fraction of focal adhesions with TagBFP signals (i.e. forces larger than 5 pN) than with mScarlet-I signals (i.e. forces larger than 3 pN). In addition, our results demonstrate that TagBFP signals are further away from the cell edge, likely coinciding with more mature focal adhesions (Fig. 3l)^4,11,75,79^. Overall, our design of mechano-sensors enabled us to collect new data on vinculin in a short time and with a high throughput, to produce a multi-view study of the forces on vinculin with unprecedented precision

In summary, we present here a new modular approach for *in vivo* force measurement using mechano-sensors that are genetically encoded, small, compatible with a versatile readout selection, and accessible to virtually any lab^83^. We expect the calibrated coiled coils are compatible with a broader range of readouts not directly tested in this paper, including FRET-based readouts^84^. The validity of our approach is independently demonstrated on different proteins and in multiple biological systems. We envision this modular architecture of mechano-sensors to herald an explosion of *in vivo* force measurements and to truly open the gate to quantitative mechanobiology.

### Limitations of the study

In this study, we have not systematically characterized in details all the different combinations of coiled coils and readouts *in vitro*. However, the modularity of our designs and the cross-validation of measurement *in vivo* using different readouts we have performed suggest each module is likely independent from each other. It is possible that due to differences in the dissociation rates of the binding pairs, the disappearance of signal after force is released vary slightly depending on the live reporter readout used, which may potentially create a discrepancy in the measurement of force thresholds between different readouts (systematic error). Another limitation of the live reporters presented in this paper is that they rely on local enrichment of fluorescence from the cytoplasm to the subcellular structures containing the proteins under force. Therefore, it may require optimization of the expression of the fluorescent binder probes, and the parameters for image acquisition and image processing to increase the signal-to-noise ratio. Supplementary note S1 presents a mathematical model and simulations for the sensors and their readouts, and provides a detailed discussion of the biochemical and biophysical parameters to consider for best results.

## KEY RESOURCES TABLE

| REAGENT or RESOURCE | SOURCE | IDENTIFIER |
| --- | --- | --- |
| Antibodies |  |  |
| Monoclonal ANTI-FLAG M2 antibody | Millipore | F3165 |
| Anti-HA High Affinity | Roche | 3F10<br>(11867423001) |
| Lamin A Monoclonal Antibody (133A2) | Invitrogen | MA1-06101 |
| Chemicals, peptides, and recombinant proteins |  |  |
| PreScission Protease | Sigma-Aldrich | GE27-0843-01 |
| d-biotin | Avidity | Bio-500 |
| Glutathione Sepharose 4B beads | Cytiva | 17075601 |
| polystyrene beads | Spherotech | PP10-20-10 |
| Lipofectamin3000 | ThermoFisher | L3000001 |
| See Table S4 for sequences of calibrated coiled-coils | This study | N/A |
| Critical commercial assays |  |  |
| Nano-Glo live cell assay system | Promega | N2011 |
| Nano-Glo HiBiT Lytic detection system | Promega | N3030 |
| Softwell 96 HTS | Matrigen | SW96G-HTS-COL-EA |
| Experimental models: Cell lines |  |  |
| Human: U2OS | Laboratory of Christian Schlieker | N/A |
| Human: U2OS with stable integration of H2B-tagBFP-P2A-GFP1-10 | Made in the Berro lab | N/A |
| Experimental models: Organisms/strains |  |  |
| See Table S1 for the list of *S. pombe* strains | This study | N/A |
| See Table S2 for the list of *C. elegans* strains | This study | N/A |
| See Table S3 for the sequences of DNA templates to generate *C. elegans* strains | This study | N/A |
| Recombinant DNA |  |  |
| pGEX6PI-Oakley-p1 | This study | pJB297 |
| pGEX6PI-Oakley-p1-OnePlus | This study | pJB298 |
| pGEX6PI-Oakley-p1-LV-L3S | This study | pJB337 |
| pGEX6PI-Oakley-p1-TwoPlus | This study | pJB339 |
| pGEX6PI-Oakley-p1-FourPlus | This study | pJB340 |
| pmCherry-Vinculin-E883-4G-DyneinStalk-GFP11 | This study | pJB348 |
| pmCherry-Vinculin-E883-4G-Oakley-p1-LV-L3S-GFP11 | This study | pJB349 |
| pmEGFP-IAAL-E3 | This study | pJB355 |
| pmCherry-Vinculin-E883-4G-DyneinStalk-IAAL-K3 | This study | pJB357 |
| pmCherry-Vinculin-E883-4G-Oakley-p1-GFP11 | This study | pJB363 |
| pmCherry-Vinculin-E883-4G-Oakley-p1-OnePlus-GFP11 | This study | pJB364 |
| pmCherry-Vinculin-E883-4G-Oakley-p1-LV-L3S-HiBiT | This study | pJB413 |
| pmScarlet3-IAAL-E3-tPT2A-Vinculin-E883-4G-DyneinStalk-IAAL-K3 | This study | pJB417 |
| pmCherry-Vinculin-E883-4G-DyneinStalk-HiBiT | This study | pJB425 |
| pmCherry-Vinculin-E883-4G-Oakley-p1-OnePlus-HiBiT | This study | pJB427 |
| pmCherry-Vinculin-E883-4G-Oakley-p1-HiBiT | This study | pJB435 |
| pmEGFP-mScarlet3-IAAL-E3-tPT2A-Vinculin-E883-4G-DyneinStalk-IAAL-K3 | This study | pJB439 |
| pTagBFP-SspB_nano-tPT2A-Vinculin-E883-4G-Oakley-p1-LV-L3S-SsrA | This study | pJB449 |
| Software and algorithms |  |  |
| GraphPad Prism 7 | GraphPad | https://www.graphpad.com |
| ImageJ | NIH | https://imagej.net |
| MATLAB | MathWorks | https://www.mathworks.com/products/matlab.html |

## Material and Methods

### Protein purification and biotinylation

DNA sequences coding coiled coils were synthesized from IDT and subcloned into BL21 (DE3) E. coli (New England BioLabs) for protein expression. Coiled coils were expressed with GST fusion at the N-terminus and Avi-tag at the C-terminus. Cleared bacteria lysates were purified by binding to Glutathione Sepharose 4B beads (GE Healthcare). The GST tag was cleaved by PreScission Protease (Sigma-Aldrich) afterwards. Avi-tag in purified coiled coil proteins was exchanged into biotinylation buffer (25 mM HEPES, 200 mM potassium glutamate, pH 7.7) and biotinylated with 50 mM bicine buffer, 50 µg/mL BirA, pH 8.3, 10 mM ATP, 10 mM magnesium acetate, and 50 μM d-biotin (Avidity) at 4°C overnight.

### DNA handle preparation and crosslinking with protein

The DNA handle used for protein attachment in single-molecule experiments was produced via PCR, yielding a 2,260 base pair fragment. This DNA fragment was modified to include a thiol group (−SH) at one end and two digoxigenin moieties at the other. For the crosslinking process, as described previously^85,86^, DNA handle was conjugated first with DTDP at pH 5.5 and excess DTDP was removed by spin column. Then purified proteins were mixed with the DTDP-treated DNA handle at a molar ratio of 50:1 (protein to DNA). This mixture was prepared in a 100 mM phosphate buffer containing 500 mM NaCl at pH 8.5 and incubated overnight at room temperature.

### Single-molecule manipulation experiments

All single-molecule experiments were performed using dual-trap high-resolution optical tweezers^85,87,88^. In brief, the two optical traps were created by focusing two orthogonally polarized beams through a water-immersed 60× objective with a 1.2 numerical aperture (Olympus). These beams originated from a single 1,064-nm laser generated by a solid-state laser (Spectra-Physics) and were then split. One of the beams was deflected by a mirror mounted on a piezoelectrically controlled stage (Mad City Labs), which could tilt along two orthogonal axes, allowing for relative movement between the traps. The outgoing laser beams were collimated by a second water immersion objective, split, and projected onto two position-sensitive detectors (Pacific Silicon Sensor). Bead displacements were detected using back-focal plane interferometry. To prepare the DNA handle crosslinked proteins, they were incubated with anti-digoxigenin antibody-coated polystyrene beads (2.17 μm in diameter, Spherotech) for 15 minutes. This mixture was then diluted with 1 mL of phosphate-buffered saline (PBS) and introduced into the top channel of a microfluidic chamber. Meanwhile, streptavidin-coated beads (1.86 μm in diameter) were introduced into the bottom channel of the chamber. Both channels were connected to a central channel via capillary tubes, where the beads were trapped. Once a bead from each type was captured, a single protein was tethered between them by bringing the two beads close together. The tethered molecule underwent pulling and relaxation by adjusting the trap separation at a rate of 10 nm/s. The optical tweezers experiments were conducted in PBS at 23 (± 1) °C. The buffer in the central channel was supplemented with an oxygen scavenging system as described elsewhere. Data were processed using MATLAB codes, also described elsewhere^85,89^, and unfolding forces were determined from the force-extension curves.

### Steered molecular dynamics (SMD) simulation

Structures of coiled coils were predicted from AlphaFold2^90^. We rotated the coiled coil so that the vector from the N terminus to the C terminus aligned with the x-axis, ensuring a consistent initial condition for all coiled coils. We used Qiwik to generate NAMD’s configuration files. The total displacement was set to 200 Å with a margin of 15 Å for water molecules, and the salt concentration was set to 0.15mol/L. We anchored the first residue of the coiled coils and pulled the last one. The speed at which we pulled the SMD’s dummy atom was 2.5e-5 A/fs (2500 m/s), and the spring constant was 1e-2kcal/mol/Å_2_ (6.949e_-1_ pN/ Å). This unrealistically high pulling speed was set up to obtain results within a reasonable time (∼60 ns of simulated time, or ∼1 week simulation, per coiled coil).

Before running the SMD simulation, we performed a minimization for 5,000 steps, followed by annealing by increasing the temperature from 60 K to 300 K. We increased the temperature by 1-degree increment and ran 600 steps between each increment. In total, it took 14,400 steps to increase the temperature to 300 K. Finally, we equilibrated the system by simulating it for 500,000 steps. We used the result of the equilibrated system as the initial condition of the SMD simulation. We used a step size of 2 fs for all the simulations mentioned.

To measure the force threshold, we measured the minimum force required to open the coiled coil in each iteration and then took the average over all forces. We used the Potential of Mean Force as described by Park and Schulten^91^.

### Selection of insertion sites for force sensors

To choose insertion sites on a protein of interest, we typically avoid known folded domains, and preferentially choose a disordered region that has no known post-translational modification or binding partner using protein databases that are species specific (e.g. Pombase for *S. pombe* or Wormbase for *C. elegans*) or general (e.g. Uniprot). When possible, we recommend using protein structure prediction tools (e.g. AlphaFold) to check the protein structure with and without the insertion of the force sensors.

### Yeast strains and media

The *S. pombe* strains in this paper are listed in Supplementary Table S1. The strains were created through CRISPR-mediated genome editing as outlined previously^92–94^ and confirmed through sequencing of colony PCR products. Yeast cells were cultured in YE5S medium (Yeast Extract supplemented with 0.225 g/L each of uracil, adenine, lysine, histidine, and leucine) and imaged on gelatin pads made with EMM5S (Edinburgh Minimum media supplemented with 0.225 g/L each of uracil, adenine, lysine, histidine, and leucine). Before imaging, yeast cells were grown at 32 °C with continuous shaking at 200 rpm overnight, allowing them to reach the exponential growth phase with OD_595nm_ values between 0.3 and 0.5.

### Confocal microscopy for yeast cells

Live imaging of *S. pombe* cells was performed on a 25% gelatin pad at room temperature, using a Nikon TiE inverted microscope with a CSU-W1 Confocal Scanning Unit from Yokogawa Electric Corporation (Tokyo, Japan). The microscope was equipped with a CFI Plan Apo 100X/1.45NA Phase objective from Nikon. Image acquisition was performed using an iXon Ultra888 EMCCD camera from Andor (Belfast, UK). For imaging GFP, an excitation wavelength of 488nm from an argon-ion laser was used, and the fluorescence emission was filtered via a single band pass filter 510/25 in the Spectra X system. mCherry and sfCherry2 were excited from a 561nm argon-ion laser, and the fluorescence emission was filtered through a single band pass filter 575/25 in the Spectra X system. To image the entire yeast cell, 21 optical slices were collected with a thickness of 0.5 µm, followed by maximum intensity projection to generate 2D images. For image display and analysis, the Fiji distribution of ImageJ provided by the National Institutes of Health (NIH, USA) was used^95^. *S. pombe* cells were also imaged on gelatin pads on BC43 CF tabletop confocal (Andor) with 100x Plan Apochromat oil immersion objective and preset excitation/emission combinations for GFP, mCherry and sfCherry.

### TIRF microscopy

Live TIRF imaging of yeast cells was performed at room temperature using gelatin pads prepared as previously described^96,97^. Imaging was conducted using a Nikon Ti2E inverted microscope equipped with an Abbelight module (Cachan, France), equipped with a Teledyne Kinetix Scientific CMOS camera (01-KINETIX-M-C; Teledyne Photometrics, AZ, USA), 488-nm and 561-nm solid-state diode lasers, 525/50 and 600/40 single bandpass filters (for mEGFP and mScarlet-I imaging, respectively). Both channels were imaged simultaneously using a beam splitter, at 1-second intervals. Image display and analysis were performed using the Fiji distribution of ImageJ (National Institutes of Health, USA)^95,98^.

### Measurement of bioluminescence (yeast)

Yeast cells were grown to exponential phase in YE5S medium, pelleted, and resuspended in EMM5S to a final density of 1 OD_595nm_/μL. 150 μL cells were mixed with 15 μL diluted substrate (Nano-Glo Live, Promega, 1:50 dilution.) and loaded into a black 96 well plate (#675086, Greiner Bio-One) before measuring luminescence on a plate reader (BioTek synergy H1, Agilent). Luminescence signals were recorded 5 minutes after mixing cells with the substrate and normalized to the OD_595nm_ of each well. Measurements were kept in a linear range predetermined as in Fig. S3.

### Measurement of bioluminescence (mammalian cells)

U2OS cells were cultured in a 4-compartment dish (#627870, Greiner Bio-One) till ∼90% confluency. Cells were transfected with plasmids (pJB425, pJB413, pJB435, pJB427) using Lipofectamine 3000. 24 Hours after transfection, cells were removed from each compartment by trypsin digestion, pelleted and separated into three equal volumes (100 μL each). One volume was mixed with 20 μL Nano-Glo Lytic buffer containing 1:50 diluted substrate and 1:100 diluted LgBiT for luminescence measurement in a black 96-well plate <5 mins after digestion. The other volumes were added either directly into a well of a black 96-well plate, or into a well precoated with 10 μL Matrigel (Corning). Luminescence was measured 6 hours after plating by adding Nano-Glo Lytic buffer containing 1:50 diluted substrate and 1:100 diluted LgBiT. For the Matrigel coated well, 10 μL culture media was taken out before measurement to keep the volumes consistent. Raw luminescence values were normalized by cell density in each well. For luminescence measurements on mCherry-vinculin, mCherry signals were used for normalization.

To measure the forces on vinculin across stiffness gradients, U2OS cells were transfected with plasmids (pJB425, pJB413, pJB435, pJB427) using Lipofectamine 3000. 24 Hours after transfection, cells were collected by trypsin digestion, pelleted and separated into 11 equal volumes (100 μL each) before being replated onto a 96 well plate with gradient stiffness (SW96G-HTS-COL-EA, Matrigen) overnight. Luminescence was measured by adding 20 μL Nano-Glo Lytic buffer containing 1:50 diluted substrate and 1:100 diluted LgBiT. Raw luminescence values were normalized by mCherry-vinculin signals in each well.

### FRAP of vinculin live force reporter

U2OS cells were co-transfected with pJB355 and pJB357 following standard protocols with Lipofectamine 3000. Cells were imaged 24 hours after transfection. FRAP was performed on FV3000 using a 60X objective (NA 1.42). After correcting for photobleaching (ImageJ, NIH), the recovery of fluorescence was fitted with a regression curve f(t)=MF*(1-exp(−k*t)), where MF is a mobile fraction and k is a turnover rate.

### Nematode Microscopy and Data Analysis

Nematodes were cultured at 20°C on nematode growth medium plates that were seeded with OP50 bacteria. Detailed animal preparation for microscopy was published previously^99^. In brief, animals at their larval stage L4 were mounted on 10% agarose patch and paralyzed in a droplet of 10 mM Levamisole diluted in M9 medium. Images were acquired with a DMi8 inverted microscope (Leica) that is equipped with a VT-iSIM system (Biovision) and an ORCA-Flash 4.0 camera (Hamamatsu). The microscope was controlled by the MetaMorph Advanced Confocal Acquisition Software. Images were acquired with an HC PL APO 40x/1.30NA OIL CS2 objective at a 488nm laser line. Raw images were processed and analyzed in Fiji/ImageJ v2.3.0/1.53f51^95,98^. Images were acquired in single layers and then stacked into maximum projections. To capture the intensity signal along the entire length of PLM, which could not be acquired in a single field of view, multiple images along the length of the neurite were taken and stitched into a single image using the pairwise-stitching plugin with a linear blending fusion method^100^. To determine the mean fluorescence along the PLM neurite, a 5-pixel thick line was drawn along the center of the neurite and a signal intensity profile was generated by using the plot profile function and the signal intensity was averaged. To subtract background fluorescence, the same signal intensity profile was acquired by shifting the drawn line from the center of the neurite by a few pixels into the non-neuronal tissue directly contacting the neurite. The signal intensity was calculated in arbitrary units as I_mean_=I_mean_ (neurite)-I_mean_ (background).

### Plasmids and transient transfections for TEVp expressing cells

The TEV module was cloned using standard cloning techniques into the nesprin TSmod 25 in pcDNA3.1 (https://www.addgene.org/browse/sequence/244038/)^101^. The mCerulean and mVenus sequences were removed and replaced with either the 5 pN or 10 pN sensor leading to an insertion between the N-terminal actin-binding CH domain (1-485) and the C-terminal KASH SUN-binding domain (6525–6874) (Fig. 2l)

U2OS cells were cultured on coverslips and transfected using Lipofectamine 3000 reagent (ThermoFisher) according to the manufacturer’s instructions. Immunoblot protocols were executed as previously described^102^. Briefly, 10 µg of protein were resolved by 10% SDS–PAGE gels and transferred onto polyvinylidene fluoride membranes (Bio-Rad). Membranes were blocked in 4% (wt/vol) milk in PBS + 0.1% (vol/vol) Tween-20 (Sigma-Aldrich). Primary and horseradish peroxidase-conjugated secondary antibodies were diluted in blocking buffer. A ChemiDoc gel imaging system (Bio-Rad) was used to visualize chemiluminescence.

For immunofluorescence, cells were fixed with 4% PFA/PBS for 15 min, permeabilized with 0.3% Triton X-100/PBS for 3 min, and blocked with 4% BSA/PBS (Sigma-Aldrich) for 1 hour. Coverslips were incubated for one hour with primary antibodies: FLAG (Millipore Sigma F3165, 1:500), HA (Roche 11867423001, 1:500), or LaminA (Invitrogen MA1-06101, 1:500). Samples were washed 2x with PBS for 5 min, incubated with Alexa Fluor (Life Technologies) secondary antibodies for one hour, and then washed 3x with PBS and mounted with ProLong Gold Antifade reagent + DAPI (Thermo Fisher P36935). Images were acquired on an LSM 880 laser scanning confocal microscope (Zeiss) with Airyscan using a C Plan-Apochromat 63×/1.40 oil DIC M27 objective using ZEN 2.1 software (Zeiss).

### Statistical evaluation

Statistical evaluation was performed with GraphPad Prism (version 7). The dataset was first tested for normality distribution by using the D’Agostino and Pearson test to judge the use of non-vs parametric statistical tests. For formin Cdc12 and Vinculin force measurements, one-way ANOVA with Tukey’s multiple comparison tests was used with MATLAB (MathWorks, version 2021).

## Acknowledgement

We thank the Yale West Campus Imaging Core for providing resources for microscopy and Keck DNA Sequencing Facility at Yale for their assistance. We thank Dr. Chengye Feng for helping construct the *C. elegans* strains with split-GFP readouts, and Dr. Kun Zhou for *in vitro* testing of coiled coils on DNA origami force clamps. We thank Dr. Thomas Kirkland and Rahele Esmatpour Salmani of Promega for their help with the split-Nanoluc readout. The *unc-115(ky275)* strain was provided by the Caenorhabditis Genetic Center (CGC), which is funded by NIH Office of Research Infrastructure Programs (P40 OD010440). OG is supported by a Walter-Benjamin Scholarship funded by the Deutsche Forschungsgemeinschaft (DFG, German Research Foundation) Project# 465611822. SY is supported by the NIH grant R35GM131744. YZ is supported by the NIH grant R35GM131714. JB is supported by the NIH grants R21GM132661, R01GM115636, R01EB037112 and the Research Corporation for Scientific Advancement grant SA-CMC-2021-037. YR was funded by the Whitman Fellowship from the Marine Biological Laboratory for pilot force measurements in *C. elegans*.

## Declaration of interests

Julien Berro, Yuan Ren and Yongli Zhang have a pending patent application PCT/US2023/069505.

## Supplemental information

Figures S1–S11 and Table S1-S4

Note S1

## Supplementary tables

**Supplementary Table S1.**
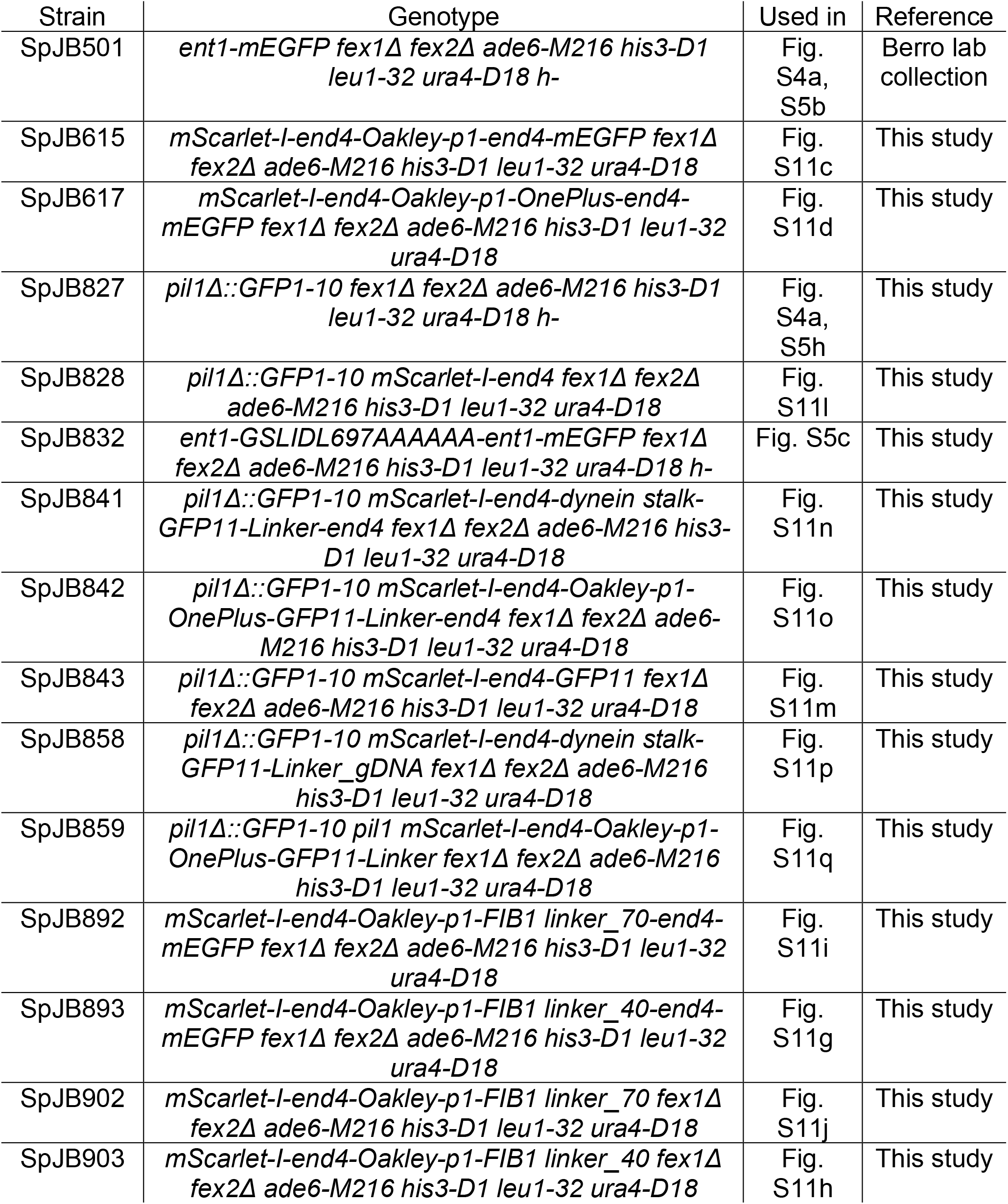
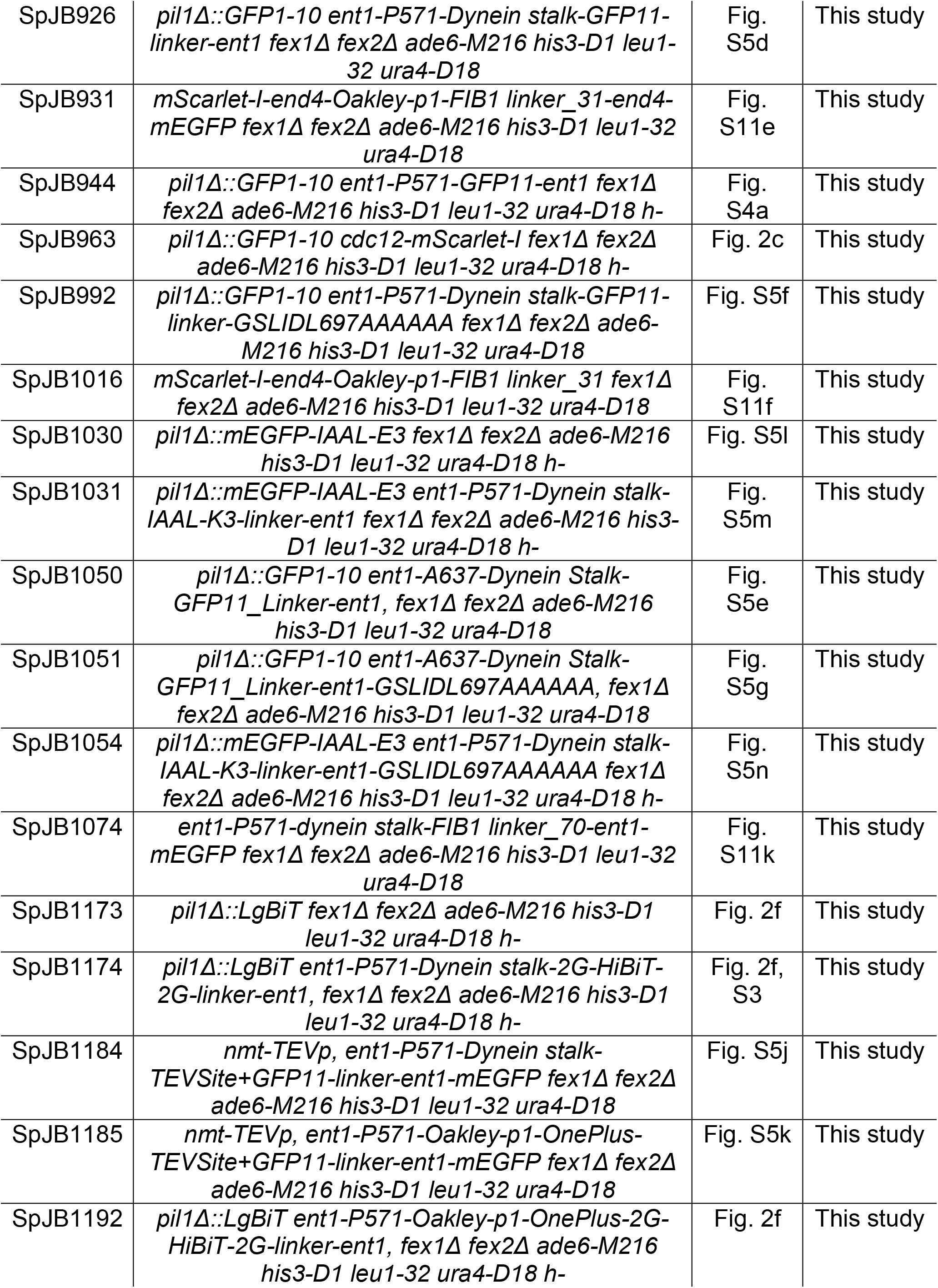
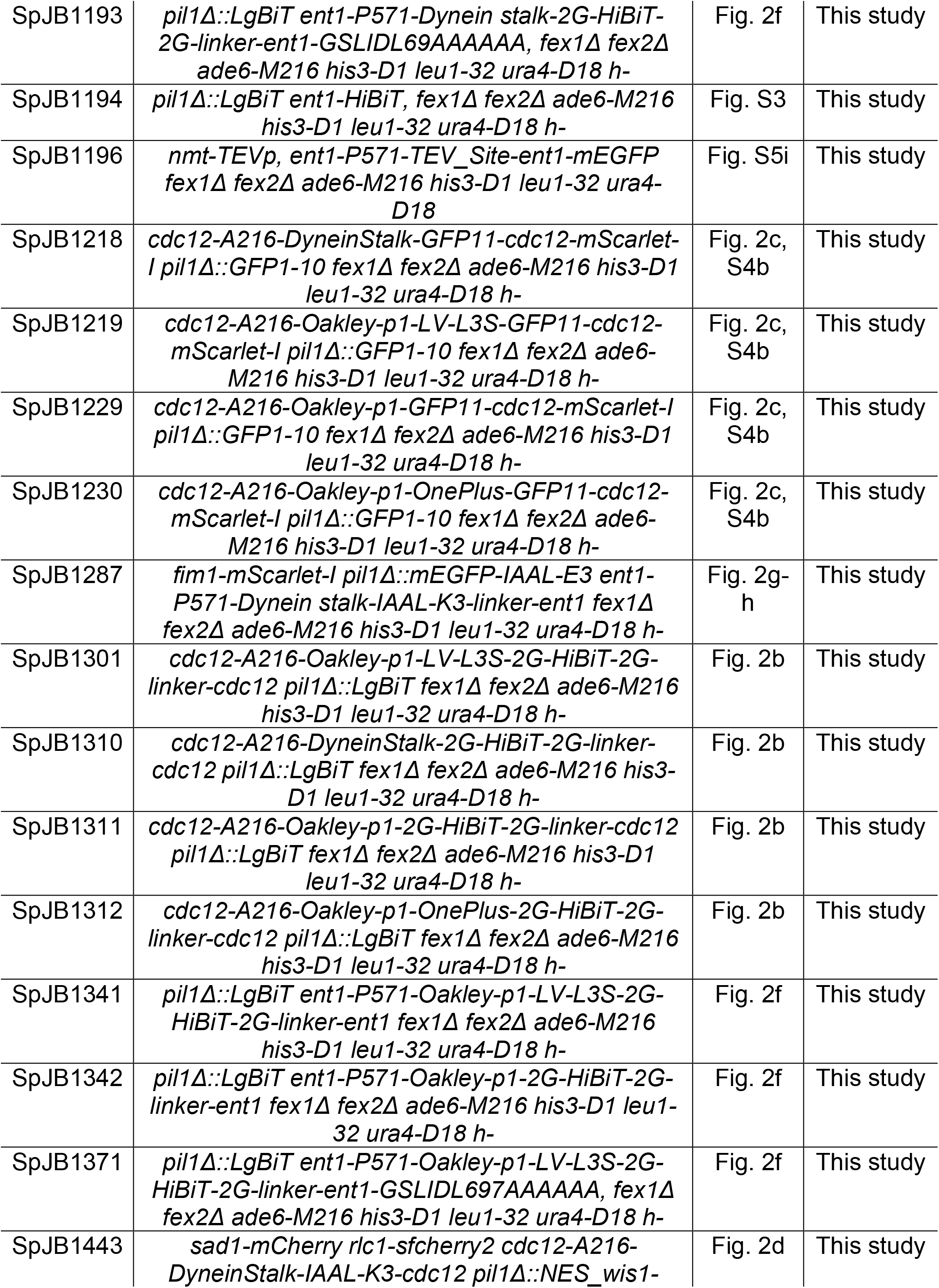
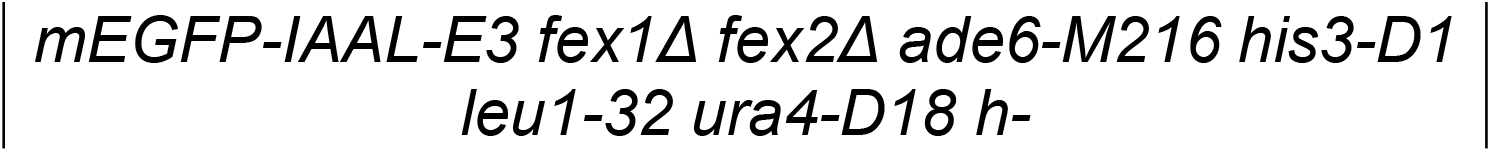
*S. pombe* strains used in this study.

| Strain | Genotype | Used in | Reference |
| --- | --- | --- | --- |
| SpJB501 | *ent1-mEGFP fex1Δ fex2Δ ade6-M216 his3-D1 leu1-32 ura4-D18 h-* | Fig. S4a, S5b | Berro lab collection |
| SpJB615 | *mScarlet-I-end4-Oakley-p1-end4-mEGFP fex1Δ fex2Δ ade6-M216 his3-D1 leu1-32 ura4-D18* | Fig. S11c | This study |
| SpJB617 | *mScarlet-I-end4-Oakley-p1-OnePlus-end4-mEGFP fex1Δ fex2Δ ade6-M216 his3-D1 leu1-32 ura4-D18* | Fig. S11d | This study |
| SpJB827 | *pil1Δ::GFP1-10 fex1Δ fex2Δ ade6-M216 his3-D1 leu1-32 ura4-D18 h-* | Fig. S4a, S5h | This study |
| SpJB828 | *pil1Δ::GFP1-10 mScarlet-I-end4 fex1Δ fex2Δ ade6-M216 his3-D1 leu1-32 ura4-D18* | Fig. S11l | This study |
| SpJB832 | *ent1-GSLIDL697AAAAAA-ent1-mEGFP fex1Δ fex2Δ ade6-M216 his3-D1 leu1-32 ura4-D18 h-* | Fig. S5c | This study |
| SpJB841 | *pil1Δ::GFP1-10 mScarlet-I-end4-dynein stalk-GFP11-Linker-end4 fex1Δ fex2Δ ade6-M216 his3-D1 leu1-32 ura4-D18* | Fig. S11n | This study |
| SpJB842 | *pil1Δ::GFP1-10 mScarlet-I-end4-Oakley-p1-OnePlus-GFP11-Linker-end4 fex1Δ fex2Δ ade6-M216 his3-D1 leu1-32 ura4-D18* | Fig. S11o | This study |
| SpJB843 | *pil1Δ::GFP1-10 mScarlet-I-end4-GFP11 fex1Δ fex2Δ ade6-M216 his3-D1 leu1-32 ura4-D18* | Fig. S11m | This study |
| SpJB858 | *pil1Δ::GFP1-10 mScarlet-I-end4-dynein stalk-GFP11-Linker_gDNA fex1Δ fex2Δ ade6-M216 his3-D1 leu1-32 ura4-D18* | Fig. S11p | This study |
| SpJB859 | *pil1Δ::GFP1-10 pil1 mScarlet-I-end4-Oakley-p1-OnePlus-GFP11-Linker fex1Δ fex2Δ ade6-M216 his3-D1 leu1-32 ura4-D18* | Fig. S11q | This study |
| SpJB892 | *mScarlet-I-end4-Oakley-p1-FIB1 linker_70-end4-mEGFP fex1Δ fex2Δ ade6-M216 his3-D1 leu1-32 ura4-D18* | Fig. S11i | This study |
| SpJB893 | *mScarlet-I-end4-Oakley-p1-FIB1 linker_40-end4-mEGFP fex1Δ fex2Δ ade6-M216 his3-D1 leu1-32 ura4-D18* | Fig. S11g | This study |
| SpJB902 | *mScarlet-I-end4-Oakley-p1-FIB1 linker_70 fex1Δ fex2Δ ade6-M216 his3-D1 leu1-32 ura4-D18* | Fig. S11j | This study |
| SpJB903 | *mScarlet-I-end4-Oakley-p1-FIB1 linker_40 fex1Δ fex2Δ ade6-M216 his3-D1 leu1-32 ura4-D18* | Fig. S11h | This study |
| SpJB926 | *pil1Δ::GFP1-10 ent1-P571-Dynein stalk-GFP11-linker-ent1 fex1Δ fex2Δ ade6-M216 his3-D1 leu1-32 ura4-D18* | Fig. S5d | This study |
| SpJB931 | *mScarlet-I-end4-Oakley-p1-FIB1 linker_31-end4-mEGFP fex1Δ fex2Δ ade6-M216 his3-D1 leu1-32 ura4-D18* | Fig. S11e | This study |
| SpJB944 | *pil1Δ::GFP1-10 ent1-P571-GFP11-ent1 fex1Δ fex2Δ ade6-M216 his3-D1 leu1-32 ura4-D18 h-* | Fig. S4a | This study |
| SpJB963 | *pil1Δ::GFP1-10 cdc12-mScarlet-I fex1Δ fex2Δ ade6-M216 his3-D1 leu1-32 ura4-D18 h-* | Fig. 2c | This study |
| SpJB992 | *pil1Δ::GFP1-10 ent1-P571-Dynein stalk-GFP11-linker-GSLIDL697AAAAAA fex1Δ fex2Δ ade6-M216 his3-D1 leu1-32 ura4-D18* | Fig. S5f | This study |
| SpJB1016 | *mScarlet-I-end4-Oakley-p1-FIB1 linker_31 fex1Δ fex2Δ ade6-M216 his3-D1 leu1-32 ura4-D18* | Fig. S11f | This study |
| SpJB1030 | *pil1Δ::mEGFP-IAAL-E3 fex1Δ fex2Δ ade6-M216 his3-D1 leu1-32 ura4-D18 h-* | Fig. S5l | This study |
| SpJB1031 | *pil1Δ::mEGFP-IAAL-E3 ent1-P571-Dynein stalk-IAAL-K3-linker-ent1 fex1Δ fex2Δ ade6-M216 his3-D1 leu1-32 ura4-D18 h-* | Fig. S5m | This study |
| SpJB1050 | *pil1Δ::GFP1-10 ent1-A637-Dynein Stalk-GFP11_Linker-ent1, fex1Δ fex2Δ ade6-M216 his3-D1 leu1-32 ura4-D18* | Fig. S5e | This study |
| SpJB1051 | *pil1Δ::GFP1-10 ent1-A637-Dynein Stalk-GFP11_Linker-ent1-GSLIDL697AAAAAA, fex1Δ fex2Δ ade6-M216 his3-D1 leu1-32 ura4-D18* | Fig. S5g | This study |
| SpJB1054 | *pil1Δ::mEGFP-IAAL-E3 ent1-P571-Dynein stalk-IAAL-K3-linker-ent1-GSLIDL697AAAAAA fex1Δ fex2Δ ade6-M216 his3-D1 leu1-32 ura4-D18 h-* | Fig. S5n | This study |
| SpJB1074 | *ent1-P571-dynein stalk-FIB1 linker_70-ent1-mEGFP fex1Δ fex2Δ ade6-M216 his3-D1 leu1-32 ura4-D18* | Fig. S11k | This study |
| SpJB1173 | *pil1Δ::LgBiT fex1Δ fex2Δ ade6-M216 his3-D1 leu1-32 ura4-D18 h-* | Fig. 2f | This study |
| SpJB1174 | *pil1Δ::LgBiT ent1-P571-Dynein stalk-2G-HiBiT-2G-linker-ent1, fex1Δ fex2Δ ade6-M216 his3-D1 leu1-32 ura4-D18 h-* | Fig. 2f, S3 | This study |
| SpJB1184 | *nmt-TEVp, ent1-P571-Dynein stalk-TEVSite+GFP11-linker-ent1-mEGFP fex1Δ fex2Δ ade6-M216 his3-D1 leu1-32 ura4-D18* | Fig. S5j | This study |
| SpJB1185 | *nmt-TEVp, ent1-P571-Oakley-p1-OnePlus-TEVSite+GFP11-linker-ent1-mEGFP fex1Δ fex2Δ ade6-M216 his3-D1 leu1-32 ura4-D18* | Fig. S5k | This study |
| SpJB1192 | *pil1Δ::LgBiT ent1-P571-Oakley-p1-OnePlus-2G-HiBiT-2G-linker-ent1, fex1Δ fex2Δ ade6-M216 his3-D1 leu1-32 ura4-D18 h-* | Fig. 2f | This study |
| SpJB1193 | *pil1Δ::LgBiT ent1-P571-Dynein stalk-2G-HiBiT-2G-linker-ent1-GSLIDL69AAAAAA, fex1Δ fex2Δ ade6-M216 his3-D1 leu1-32 ura4-D18 h-* | Fig. 2f | This study |
| SpJB1194 | *pil1Δ::LgBiT ent1-HiBiT, fex1Δ fex2Δ ade6-M216 his3-D1 leu1-32 ura4-D18 h-* | Fig. S3 | This study |
| SpJB1196 | *nmt-TEVp, ent1-P571-TEV_Site-ent1-mEGFP fex1Δ fex2Δ ade6-M216 his3-D1 leu1-32 ura4-D18* | Fig. S5i | This study |
| SpJB1218 | *cdc12-A216-DyneinStalk-GFP11-cdc12-mScarlet-I pil1Δ::GFP1-10 fex1Δ fex2Δ ade6-M216 his3-D1 leu1-32 ura4-D18 h-* | Fig. 2c, S4b | This study |
| SpJB1219 | *cdc12-A216-Oakley-p1-LV-L3S-GFP11-cdc12-mScarlet-I pil1Δ::GFP1-10 fex1Δ fex2Δ ade6-M216 his3-D1 leu1-32 ura4-D18 h-* | Fig. 2c, S4b | This study |
| SpJB1229 | *cdc12-A216-Oakley-p1-GFP11-cdc12-mScarlet-I pil1Δ::GFP1-10 fex1Δ fex2Δ ade6-M216 his3-D1 leu1-32 ura4-D18 h-* | Fig. 2c, S4b | This study |
| SpJB1230 | *cdc12-A216-Oakley-p1-OnePlus-GFP11-cdc12-mScarlet-I pil1Δ::GFP1-10 fex1Δ fex2Δ ade6-M216 his3-D1 leu1-32 ura4-D18 h-* | Fig. 2c, S4b | This study |
| SpJB1287 | *fim1-mScarlet-I pil1Δ::mEGFP-IAAL-E3 ent1-P571-Dynein stalk-IAAL-K3-linker-ent1 fex1Δ fex2Δ ade6-M216 his3-D1 leu1-32 ura4-D18 h-* | Fig. 2g-h | This study |
| SpJB1301 | *cdc12-A216-Oakley-p1-LV-L3S-2G-HiBiT-2G-linker-cdc12 pil1Δ::LgBiT fex1Δ fex2Δ ade6-M216 his3-D1 leu1-32 ura4-D18 h-* | Fig. 2b | This study |
| SpJB1310 | *cdc12-A216-DyneinStalk-2G-HiBiT-2G-linker-cdc12 pil1Δ::LgBiT fex1Δ fex2Δ ade6-M216 his3-D1 leu1-32 ura4-D18 h-* | Fig. 2b | This study |
| SpJB1311 | *cdc12-A216-Oakley-p1-2G-HiBiT-2G-linker-cdc12 pil1Δ::LgBiT fex1Δ fex2Δ ade6-M216 his3-D1 leu1-32 ura4-D18 h-* | Fig. 2b | This study |
| SpJB1312 | *cdc12-A216-Oakley-p1-OnePlus-2G-HiBiT-2G-linker-cdc12 pil1Δ::LgBiT fex1Δ fex2Δ ade6-M216 his3-D1 leu1-32 ura4-D18 h-* | Fig. 2b | This study |
| SpJB1341 | *pil1Δ::LgBiT ent1-P571-Oakley-p1-LV-L3S-2G-HiBiT-2G-linker-ent1 fex1Δ fex2Δ ade6-M216 his3-D1 leu1-32 ura4-D18 h-* | Fig. 2f | This study |
| SpJB1342 | *pil1Δ::LgBiT ent1-P571-Oakley-p1-2G-HiBiT-2G-linker-ent1 fex1Δ fex2Δ ade6-M216 his3-D1 leu1-32 ura4-D18 h-* | Fig. 2f | This study |
| SpJB1371 | *pil1Δ::LgBiT ent1-P571-Oakley-p1-LV-L3S-2G-HiBiT-2G-linker-ent1-GSLIDL697AAAAAA, fex1Δ fex2Δ ade6-M216 his3-D1 leu1-32 ura4-D18 h-* | Fig. 2f | This study |
| SpJB1443 | *sad1-mCherry rlc1-sfcherry2 cdc12-A216-DyneinStalk-IAAL-K3-cdc12 pil1Δ::NES_wis1-* | Fig. 2d | This study |

**Supplementary Table S2.**
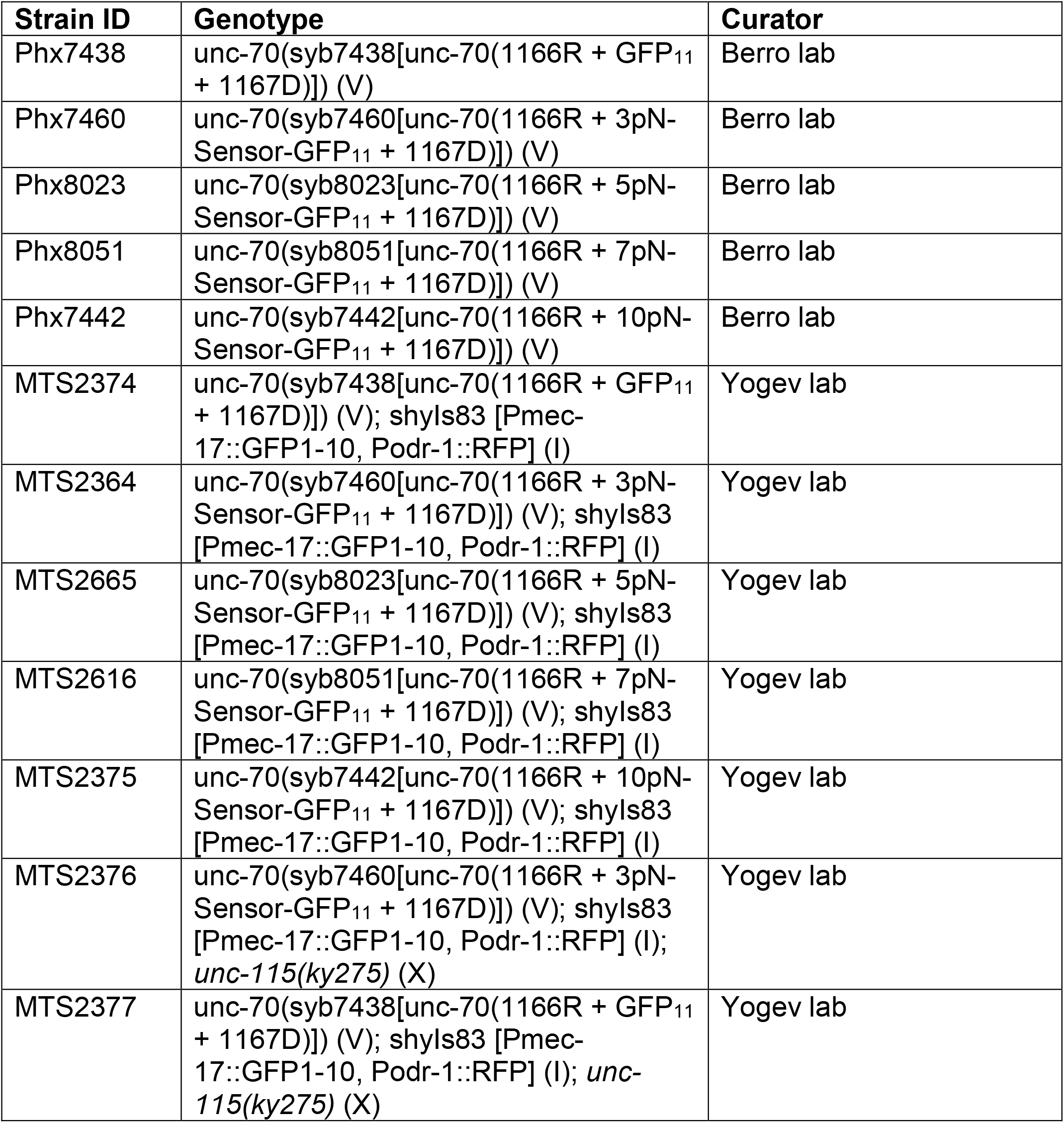
Nematode strain collection.

Nematode strain collection
| Strain ID | Genotype | Curator |
| --- | --- | --- |
| Phx7438 | unc-70(syb7438[unc-70(1166R + GFP<sub>11</sub> + 1167D)]) (V) | Berro lab |
| Phx7460 | unc-70(syb7460[unc-70(1166R + 3pN-Sensor-GFP<sub>11</sub> + 1167D)]) (V) | Berro lab |
| Phx8023 | unc-70(syb8023[unc-70(1166R + 5pN-Sensor-GFP<sub>11</sub> + 1167D)]) (V) | Berro lab |
| Phx8051 | unc-70(syb8051[unc-70(1166R + 7pN-Sensor-GFP<sub>11</sub> + 1167D)]) (V) | Berro lab |
| Phx7442 | unc-70(syb7442[unc-70(1166R + 10pN-Sensor-GFP<sub>11</sub> + 1167D)]) (V) | Berro lab |
| MTS2374 | unc-70(syb7438[unc-70(1166R + GFP<sub>11</sub> + 1167D)]) (V); shyls83 [Pmec-17::GFP1-10, Podr-1::RFP] (I) | Yogev lab |
| MTS2364 | unc-70(syb7460[unc-70(1166R + 3pN-Sensor-GFP<sub>11</sub> + 1167D)]) (V); shyls83 [Pmec-17::GFP1-10, Podr-1::RFP] (I) | Yogev lab |
| MTS2665 | unc-70(syb8023[unc-70(1166R + 5pN-Sensor-GFP<sub>11</sub> + 1167D)]) (V); shyls83 [Pmec-17::GFP1-10, Podr-1::RFP] (I) | Yogev lab |
| MTS2616 | unc-70(syb8051[unc-70(1166R + 7pN-Sensor-GFP<sub>11</sub> + 1167D)]) (V); shyls83 [Pmec-17::GFP1-10, Podr-1::RFP] (I) | Yogev lab |
| MTS2375 | unc-70(syb7442[unc-70(1166R + 10pN-Sensor-GFP<sub>11</sub> + 1167D)]) (V); shyls83 [Pmec-17::GFP1-10, Podr-1::RFP] (I) | Yogev lab |
| MTS2376 | unc-70(syb7460[unc-70(1166R + 3pN-Sensor-GFP<sub>11</sub> + 1167D)]) (V); shyls83 [Pmec-17::GFP1-10, Podr-1::RFP] (I); *unc-115(ky275)* (X) | Yogev lab |
| MTS2377 | unc-70(syb7438[unc-70(1166R + GFP<sub>11</sub> + 1167D)]) (V); shyls83 [Pmec-17::GFP1-10, Podr-1::RFP] (I); *unc-115(ky275)* (X) | Yogev lab |

**Supplementary Table S3.**
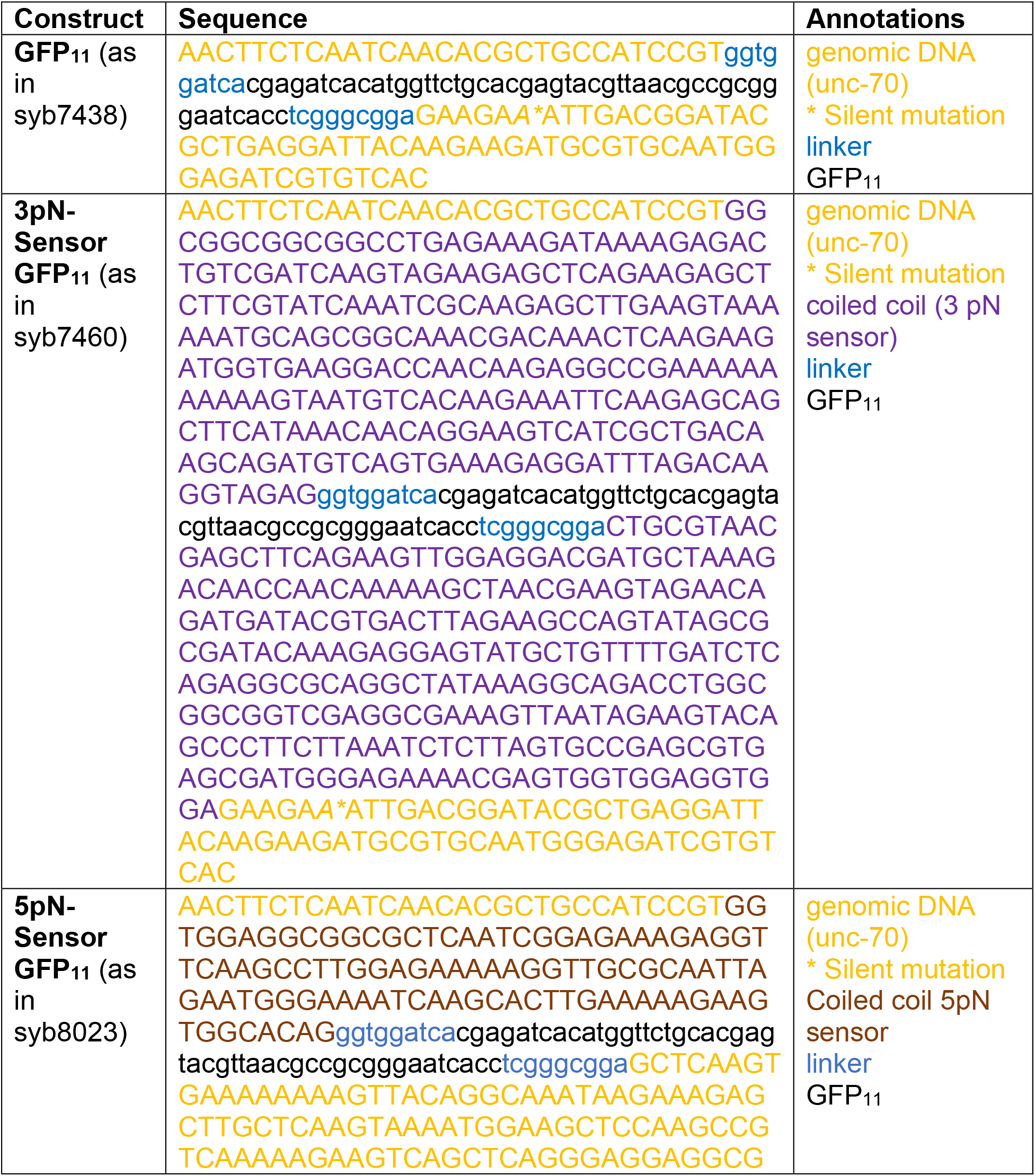
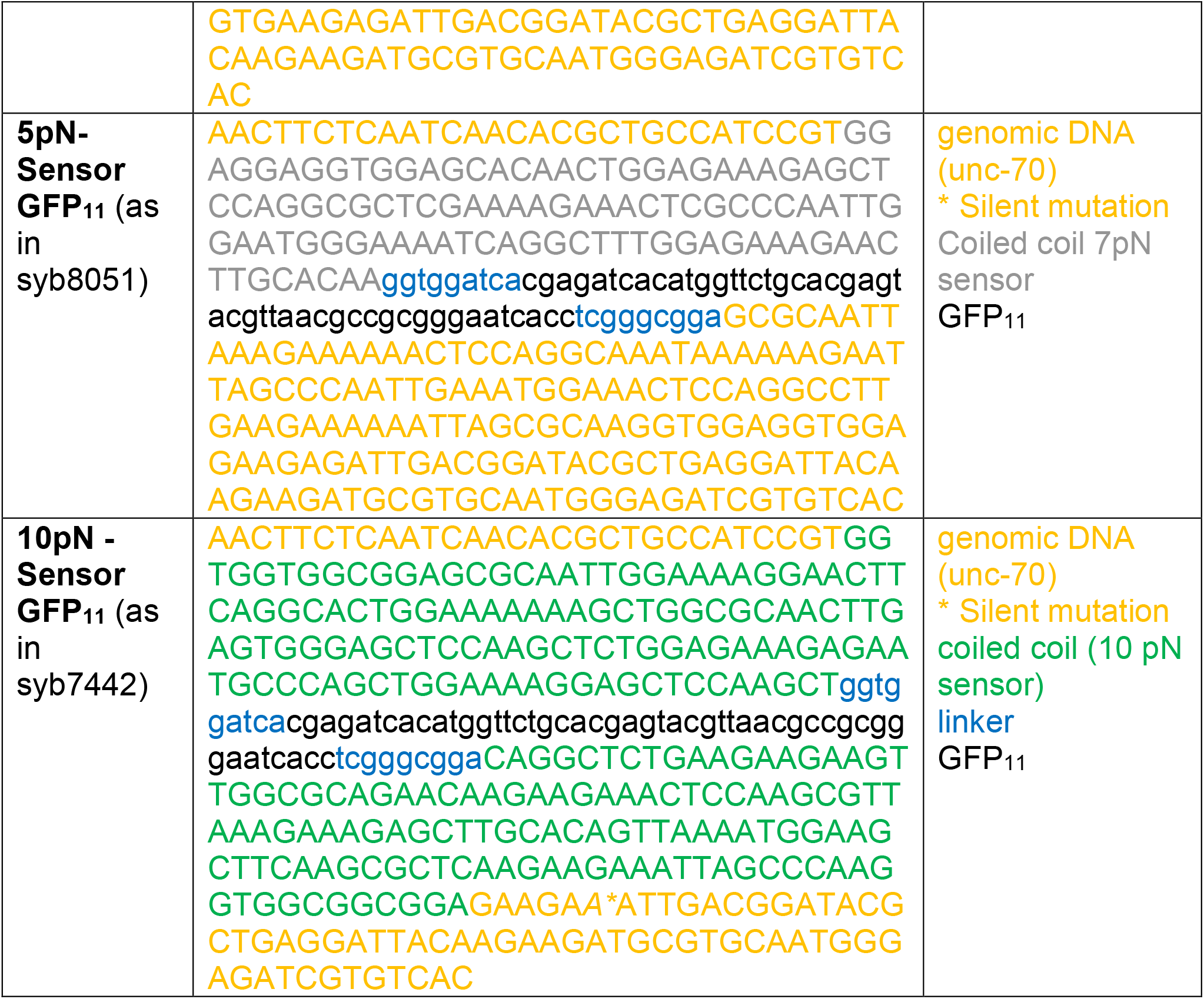
DNA templates for nematode strains. The following DNA sequences served as templates generate the alleles syb7438, syb7460, syb8023, syb8051 and syb7442, which are endogenous insertions of the force sensors into unc-70/ß-spectrin. Strains were ordered from SunyBiotech.

**Supplementary Table S4.**
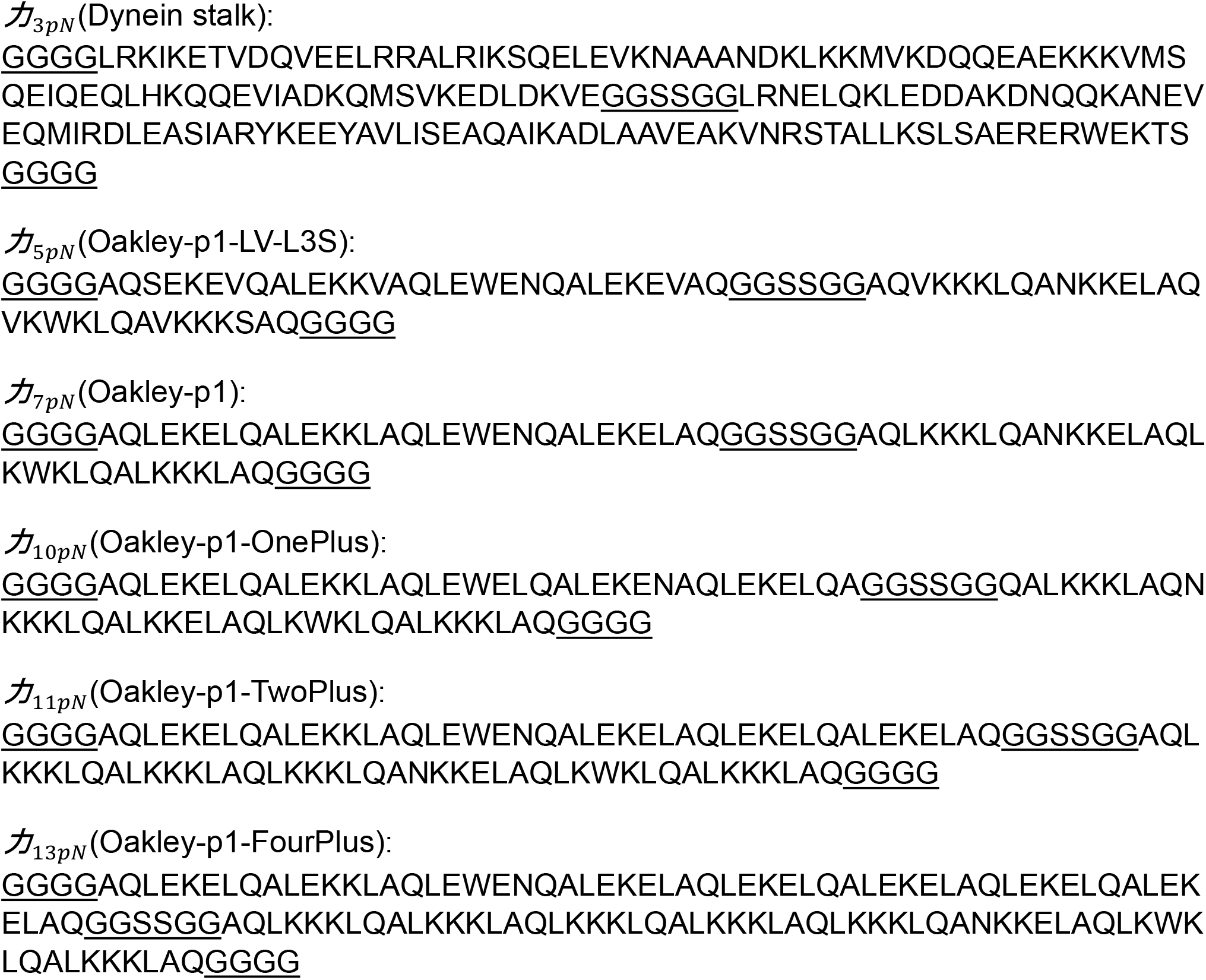
Sequences of calibrated coiled coils. Linkers are underlined.

## Supplementary figures

**Figure S1.**
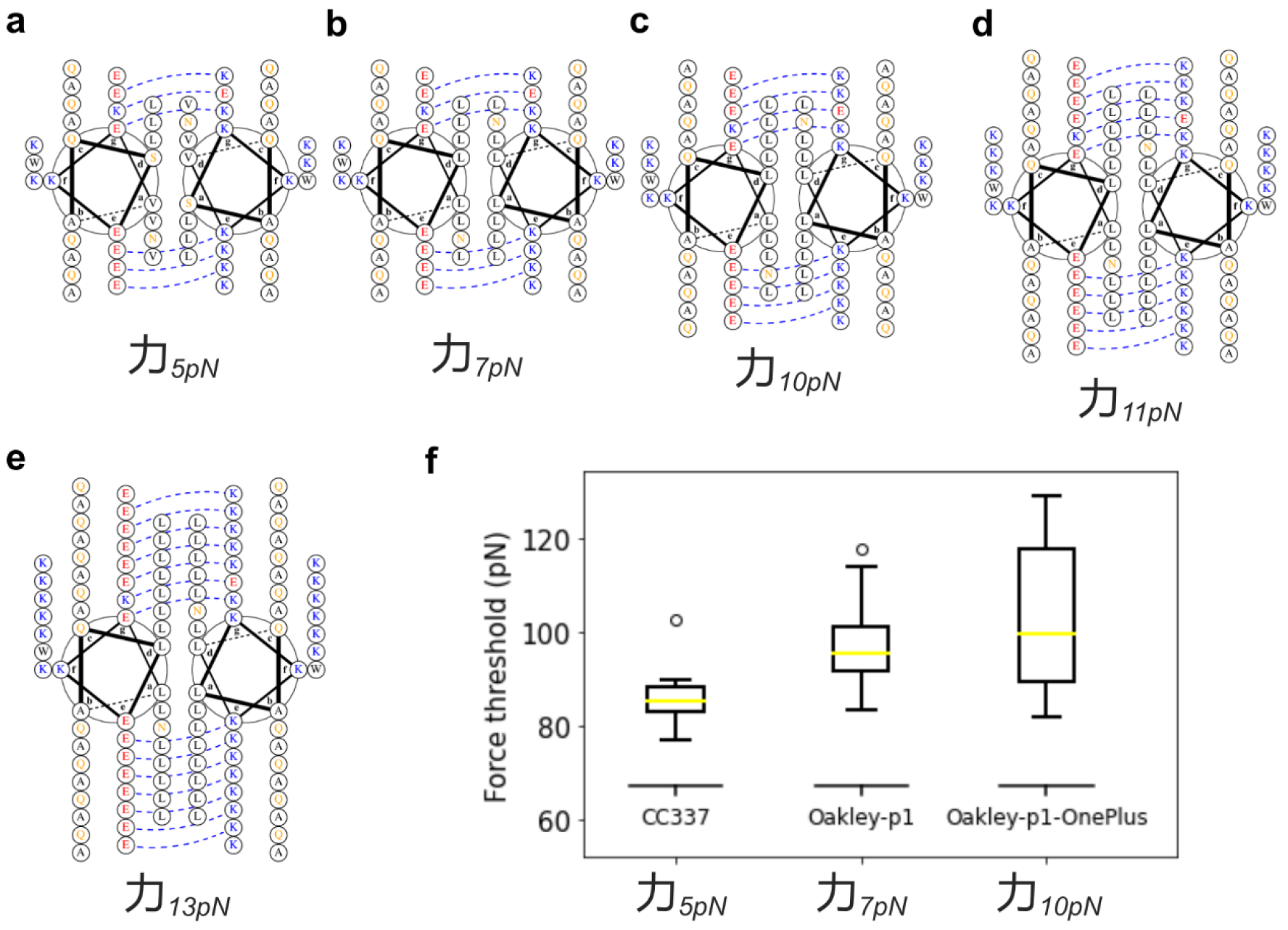
Design of heterodimeric anti-parallel coiled coils. **a-e**, The sequences of 力_5*pN*_, 力_7*pN*_, 力_10*pN*_, 力_11*pN*_, 力_13*pN*_ are shown as helical wheels (https://grigoryanlab.org/drawcoil/). E/K electrostatic charges are paired by dotted lines. **f**, Unfolding force thresholds of coiled coils from simulation. MD simulation can be used to guide coiled coil design but cannot predict the exact values of the force thresholds. See materials and methods for details.

**Figure S2.**
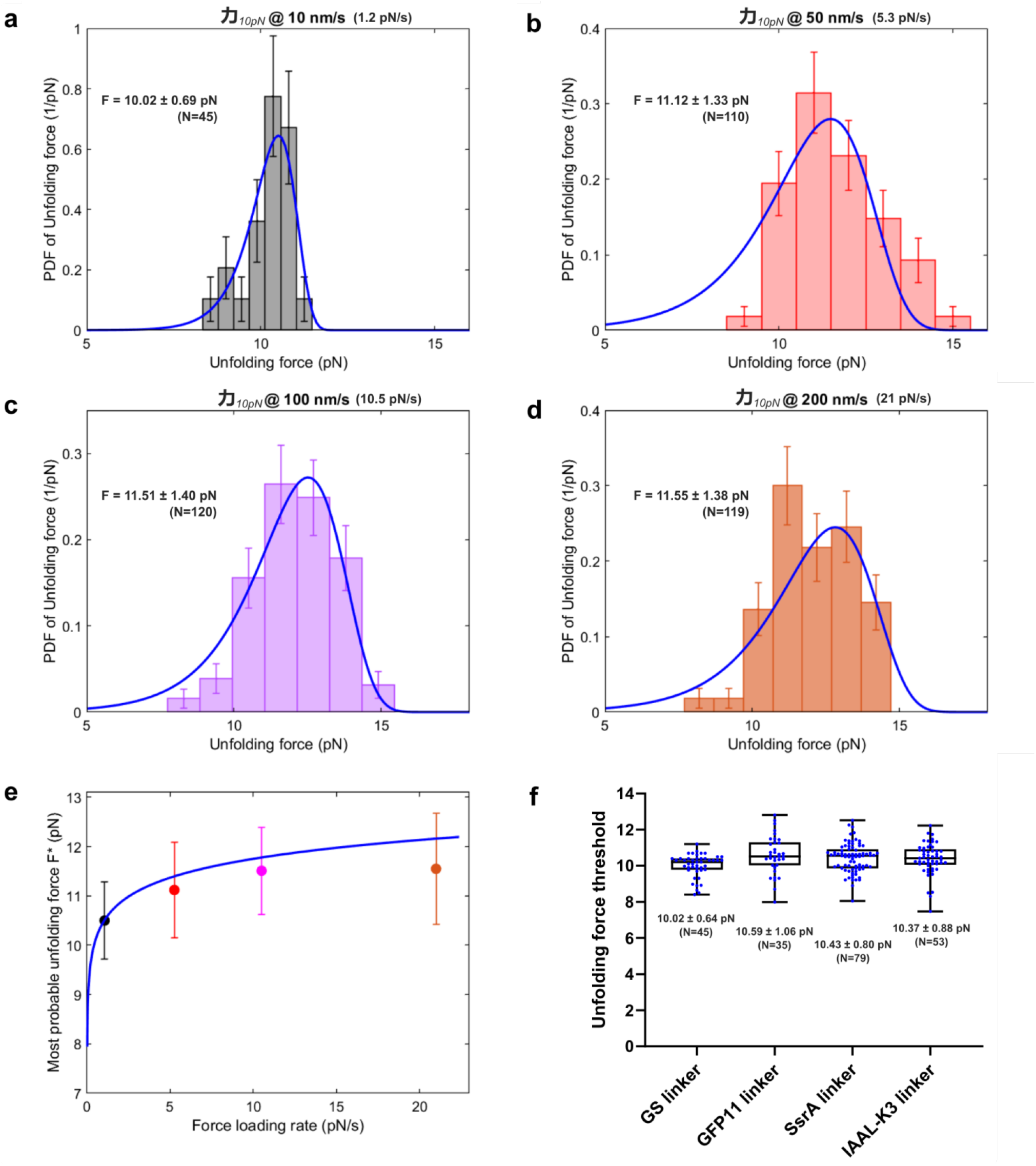
Changing the force loading rate or the readout linker have negligible influence on the coiled-coil unfolding force threshold. **a-d**, Probability distribution function (PDF) of 力_10*pN*_ pulled at different speeds from 10 nm/s to 200 nm/s. The average unfolding force F is listed as mean ± SD. **e**, The most probable unfolding force F* as a function of the force loading rate was calculated as described previously (blue line)^27^, and the calibrated unfolding forces from **a-d** are listed (colored dots). Given the approximate force loading rate of 2.3 pN/s *in vivo* and the force loading rate of ∼1.2 pN/s in optical tweezer experiments^14,15,87,89^, the difference in the unfolding force of 力_10*pN*_ reported *in vitro* and *in vivo* is less than 1 pN. The difference in the unfolding force of 力_10*pN*_ between 1 pN/s and 20 pN/s is less than 2 pN. **f**, Changing the linker of 力_10*pN*_ from a GS linker (GGSSGG) to GFP11 linker (GGRDHMVLHEYVNAAGITGG), SsrA linker (AANDENYALAA) or IAAL-K3 linker (KIAALKEKIAALKEKIAALKE) does not significantly change the unfolding force threshold. Each dot represents a pulling event.

**Figure S3.**
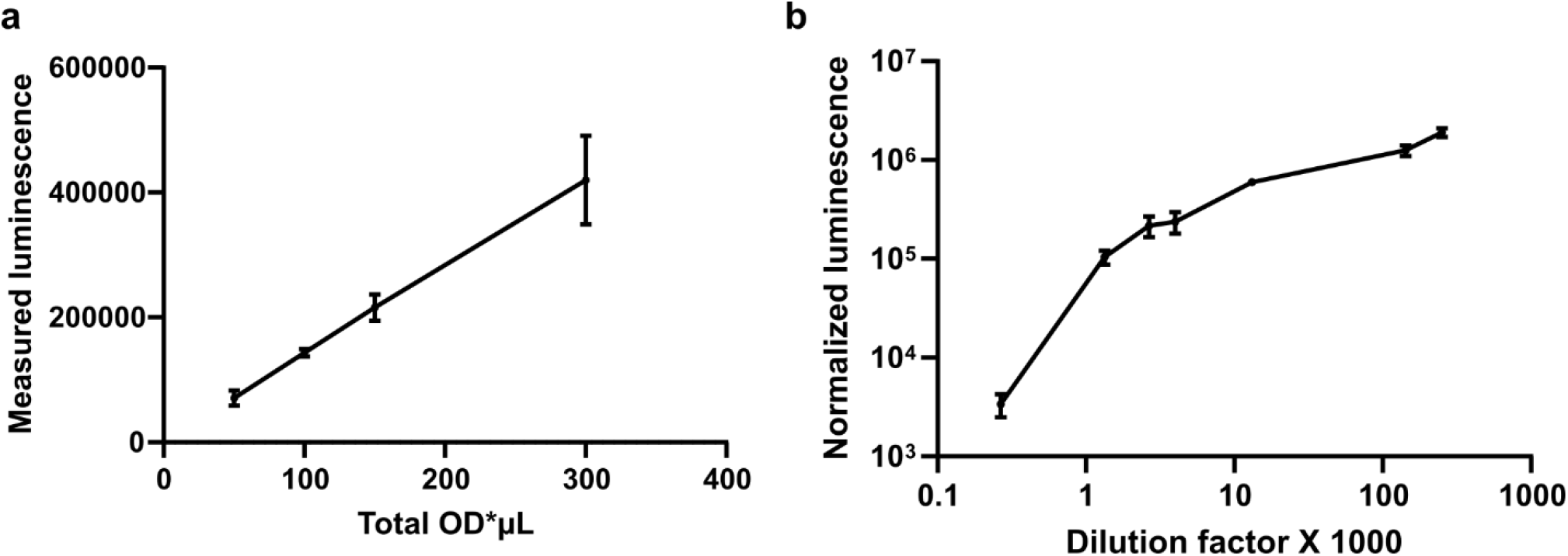
Titration of cells and substrates for luminescence measurement. **a**, Correlation between the total luminescence of fission yeast cells and the number of cells, represented by total OD*μL (1 OD*μL corresponds to 10^4^ cells). Substrates (Nano-Glo Live) were diluted 1:11. **b**, Correlation between normalized luminescence from fission yeast cells and the dilution of substrates. 100 OD*μL cells were used for each measurement, and the luminescence readings were normalized by the OD measurements on the plate reader. Error bars are SEMs. The total number of cells and the dilution of substrates were kept constant for strains expressing LgBiT and coiled-coil force sensors with HiBiT linker.

**Figure S4.**
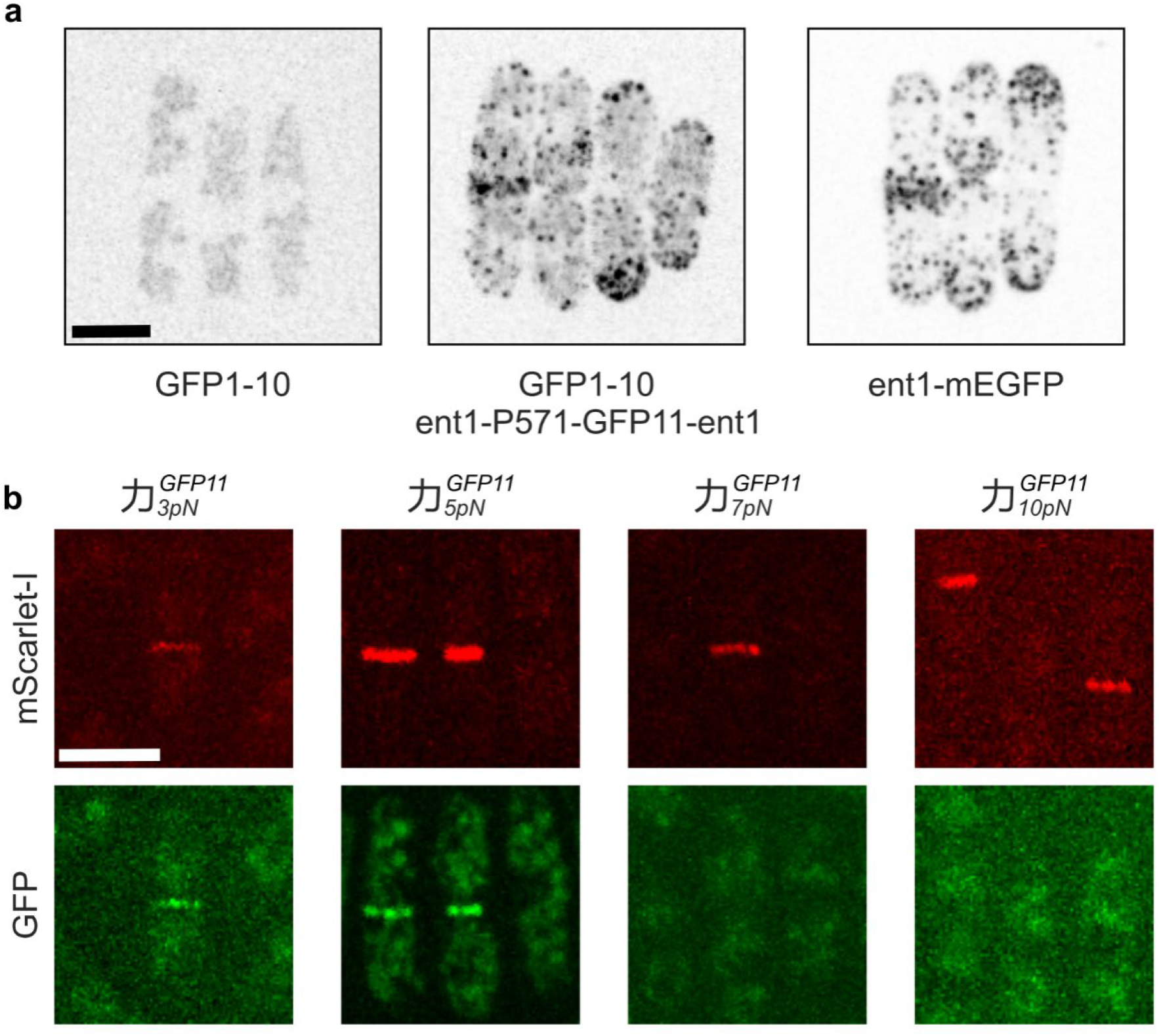
Force measurement with the split-GFP readout in fission yeast. **a**, Fission yeast cells expressing GFP1-10 alone have weak fluorescent signals (left image). Co-expression of GFP1-10 and GFP11 (inserted after P571 in Ent1, middle image) led to GFP reconstitution and the appearance of fluorescent signals with localization pattern similar to Ent1-mEGFP (right image) but with decreased intensity. Images are maximum projections of whole cell z-stacks acquired and displayed with the same settings. **b**, Quantification of forces on Cdc12 with the split-GFP readout. In cells expressing GFP1-10, endogenous Cdc12 was tagged with mScarlet-I for visualization. Force sensors (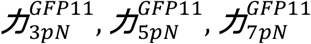 *and*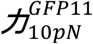) were inserted into Cdc12 after A216, and the GFP signals were normalized with respect to mScarlet-I signals to quantify force on Cdc12 (Fig. 2b). Scale bars, 5 μm.

**Figure S5.**
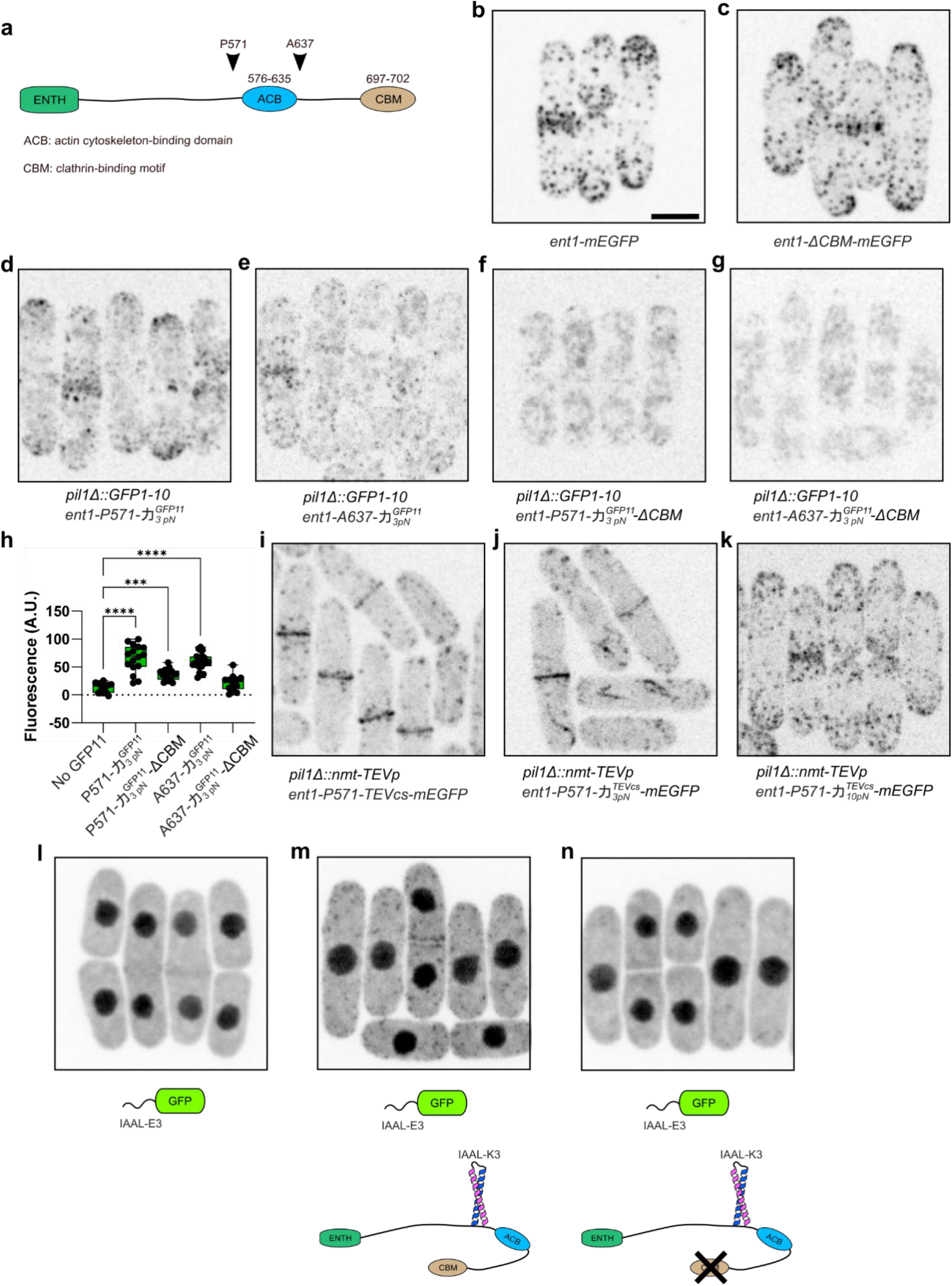
Deletion of the clathrin-binding motif (CBM) reduces the force on Ent1 both before and after the ACB domain. **a**, Schematic of Ent1 showing the insertion sites of force sensors before (P571) and after (A637) the ACB domain. Not drawn to scale. **b**, The subcellular localization of Ent1 is visualized by GFP tagging. Deletion of CBM does not change the endocytic localization of Ent1. **d-g**, In fission yeast cells expressing GFP1-10 from the pil1 locus, 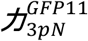 was inserted into Ent1 after P571 (**d**) or after A637 (**e**). Fluorescence signals could be detected in WT cells (**d** and **e**) but not in cells where the CBM was deleted (**f** and **g**). **h**, Quantification of fluorescence from cells containing 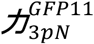 or no GFP11. ***, p<0.001. ****, p<0.0001. One-way ANOVA. **i**, The cleavage sequence TEVcs was inserted into fluorescently tagged Ent1 after P571, and TEVp expression was induced with the nmt promoter at the pil1 locus. The cleaved C-terminus of Ent1 localizes to cytokinetic F-actin rings instead of endocytic sites. **j**, The insertion of 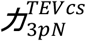 into Ent1 in TEVp-expressing cells resulted in the cleavage of Ent1, indicating force >3 pN after P571. **k**, The insertion of 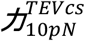 into Ent1 in TEVp-expressing cells did not cause Ent1 cleavage, indicating force <10 pN after P571. **l-n**, mEGFP-IAAL-E3 binder is enriched at endocytic sites in cells where 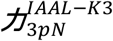 is inserted into Ent1 (**m**), indicating that force on Ent1 is above 3 pN in wild type cells. The deletion of CBM reduces force on Ent1 and led to the disappearance of locally enriched fluorescent signals (**n**). Images are maximum projections of whole cell z-stacks acquired and displayed with the same settings. Schematics below each image indicate the genotype of each strain. Scale bar, 5 μm.

**Figure S6.**
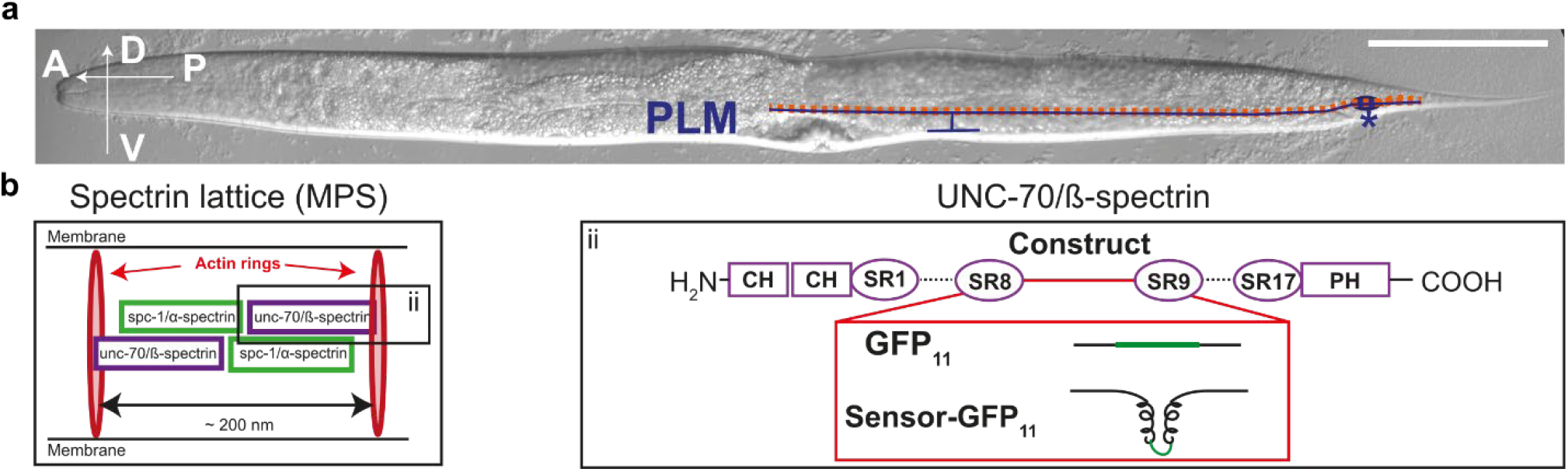
Insertion of force sensors into UNC-70 in *C. elegans*. **a**. Position of the sensory PLM neuron (blue) is illustrated in a DIC image of a nematode. Coordinates on the left show the anterior-posterior (A, P) and ventral-dorsal (V,D) axis. The PLM cell body (*) is located close to the tail of the nematode where it extends a short neurite posterior towards the tail and a long neurite anterior towards the center of the animal. Along this anterior neurite, PLM forms a short branch to innervate interneurons in the ventral nerve cord. The red dashed line along the entire length of the neurites indicates the analyzed region for Fig. 2j-k. Scale bar, 100 μm. **b**. Left panel: Spectrins form the central building block of the membrane-associated periodic skeleton (MPS), which consists of actin rings that are interspaced by spectrin tetramers and form a periodic lattice below the plasma membrane throughout the entire length of the axon. Each spectrin tetramer spaces a length of approximately 200 nm and consists of two heterodimers of α- (SPC-1 in nematodes) and ß-spectrin (UNC-70 in nematodes) subunits, which assemble head-to-head. Right panel: Protein domain organization in UNC-70/ß-spectrin with location of the tension probe insertion. A tension sensor (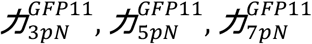 *or*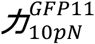) or GFP11 alone (control) were inserted using CRISPR/Cas9 into the genomic unc-70 locus at the position that encodes for the linker region between spectrin repeats 8 and 9 (after R1167). CH: Calponin Homology Domain, SR: Spectrin-like Repeats, PH: Pleckstrin Homology Domain.

**Figure S7.**
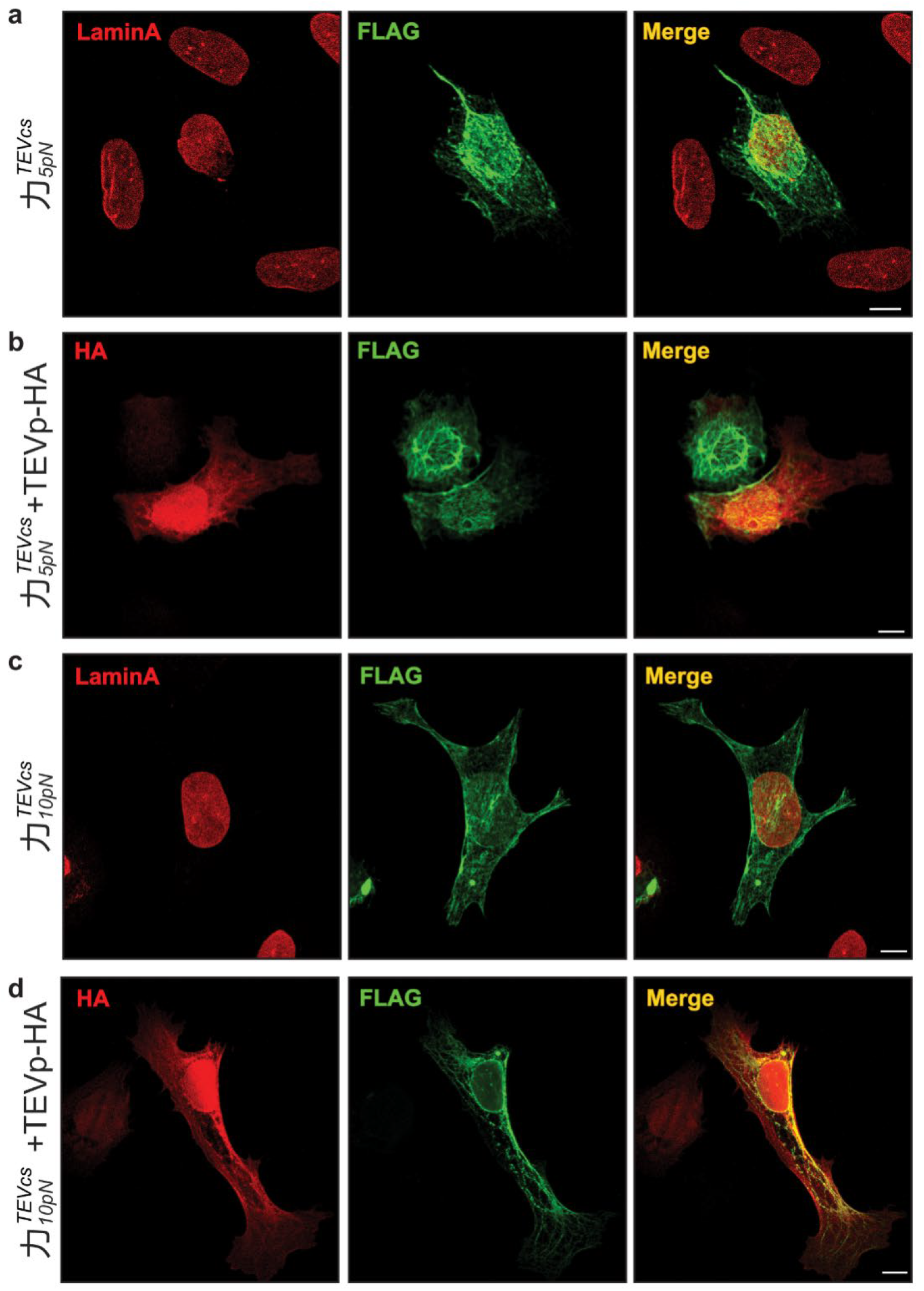
Force measurements on mini-Nesprin-2G with the TEV cleavage readout. **a-d**, Representative immunofluorescence images of U2OS cells expressing the mini-Nesprin-2G protein with the indicated force sensors 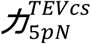(a, b) and 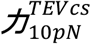(c, d) either without (a, c) or with (b, d) TEVp-HA co-expressed. Scale bar, 10 μm.

**Figure S8.**
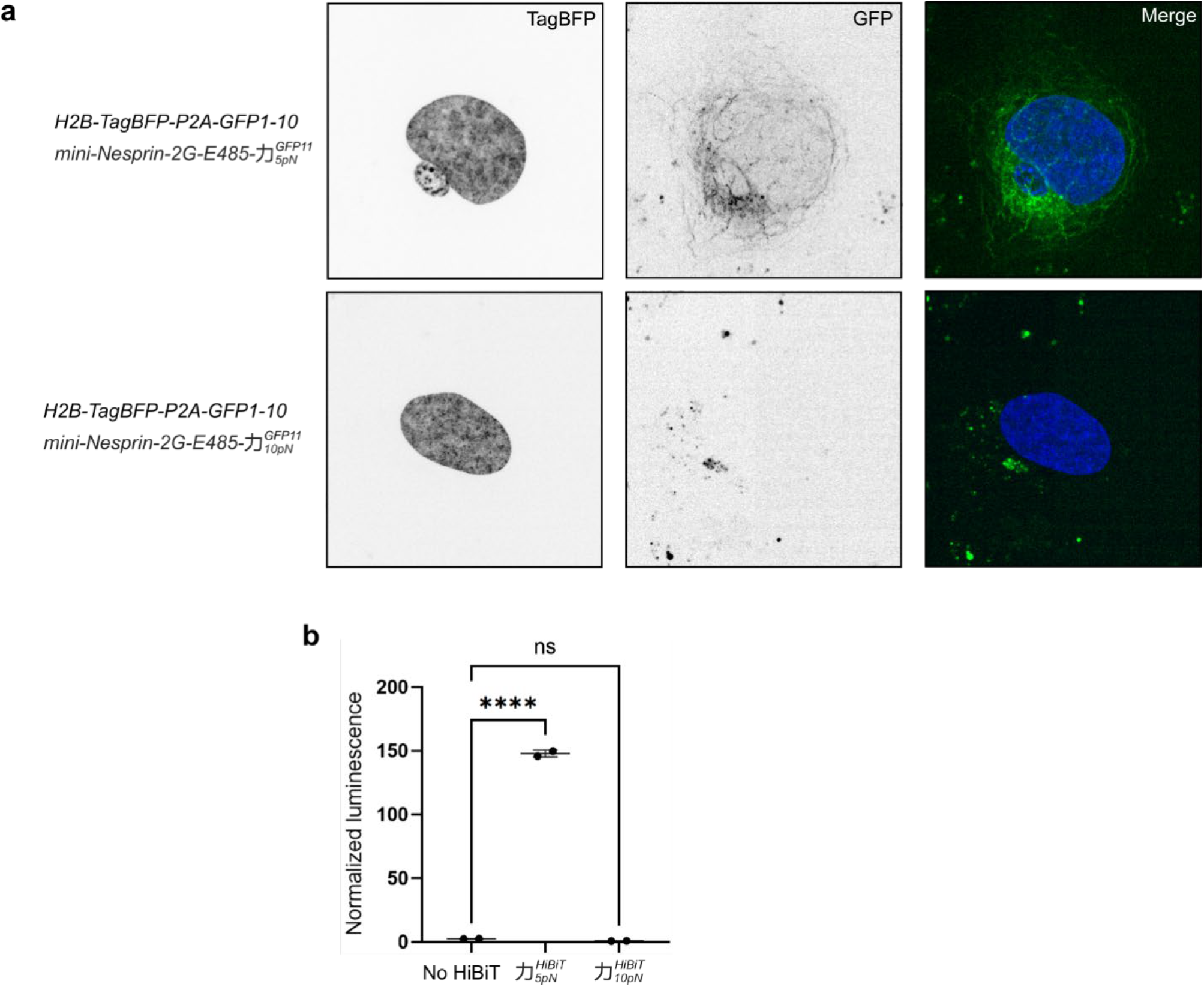
Force measurements on mini-Nesprin-2G with the split-GFP and split-NanoLuc readouts. **a**, In U2OS cells expressing H2B-TagBFP (expression marker) and GFP1-10, 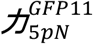 or 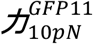 were inserted into mini-Nesprin-2G to detect forces. GFP fluorescence could be seen from cells harboring 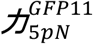 but not 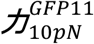, indicating a peak force between 5 pN and 10 pN. **b**, Luminescence from U2OS cells containing 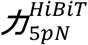 or 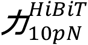 in mini-Nesprin-2G were measured and normalized to cell numbers. Significant luminescence could be detected in cells with 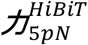 but not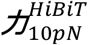. This suggests a peak force between 5 pN and 10 pN. One-way ANOVA. ****, p<0.0001. Each dot represents one independent experiment with >500 cells.

**S9.**
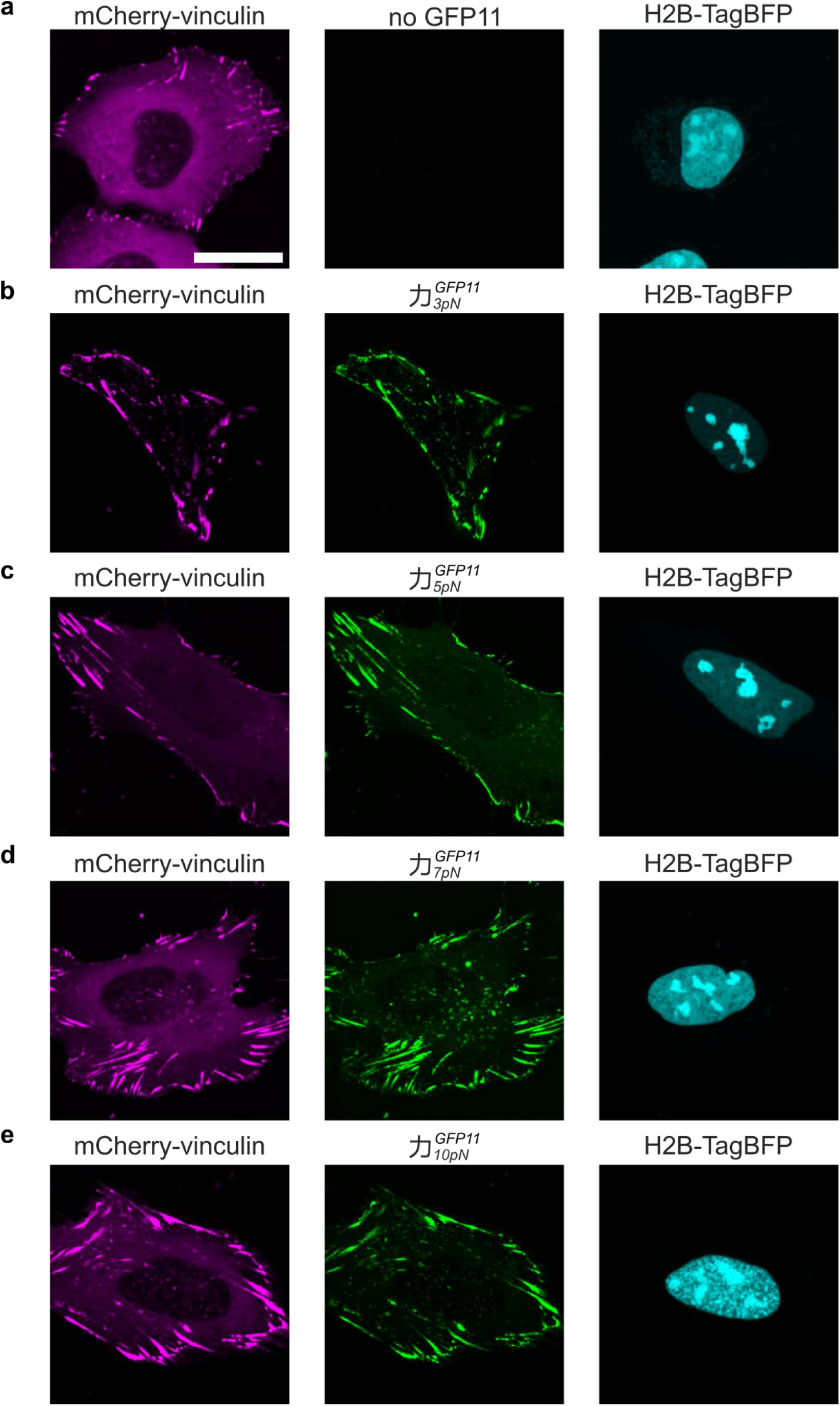
Force measurement on vinculin with the split-GFP readout. **a**, U2OS cells expressing mCherry-vinculin and H2B-TagBFP-P2A-GFP1-10 (H2B-TagBFP signal was used as an equimolar expression marker for GFP1-10) were imaged on a confocal microscope. No GFP signal was detected in cells expressing GFP1-10 but lack GFP11. **b-e**, U2OS cells expressing mCherry-vinculin with different force sensors (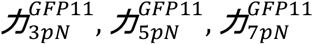 *or* 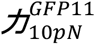) inserted after E883, and expressing H2B-TagBFP-P2A-GFP1-10 from a co-transfected plasmid. The appearance of GFP signals indicate forces above the unfolding threshold of each coiled coil sensor. The ratio between GFP and mCherry fluorescence was used as an index to quantify forces on vinculin molecules in Fig. 3b. Scale bar, 20 μm.

**S10.**
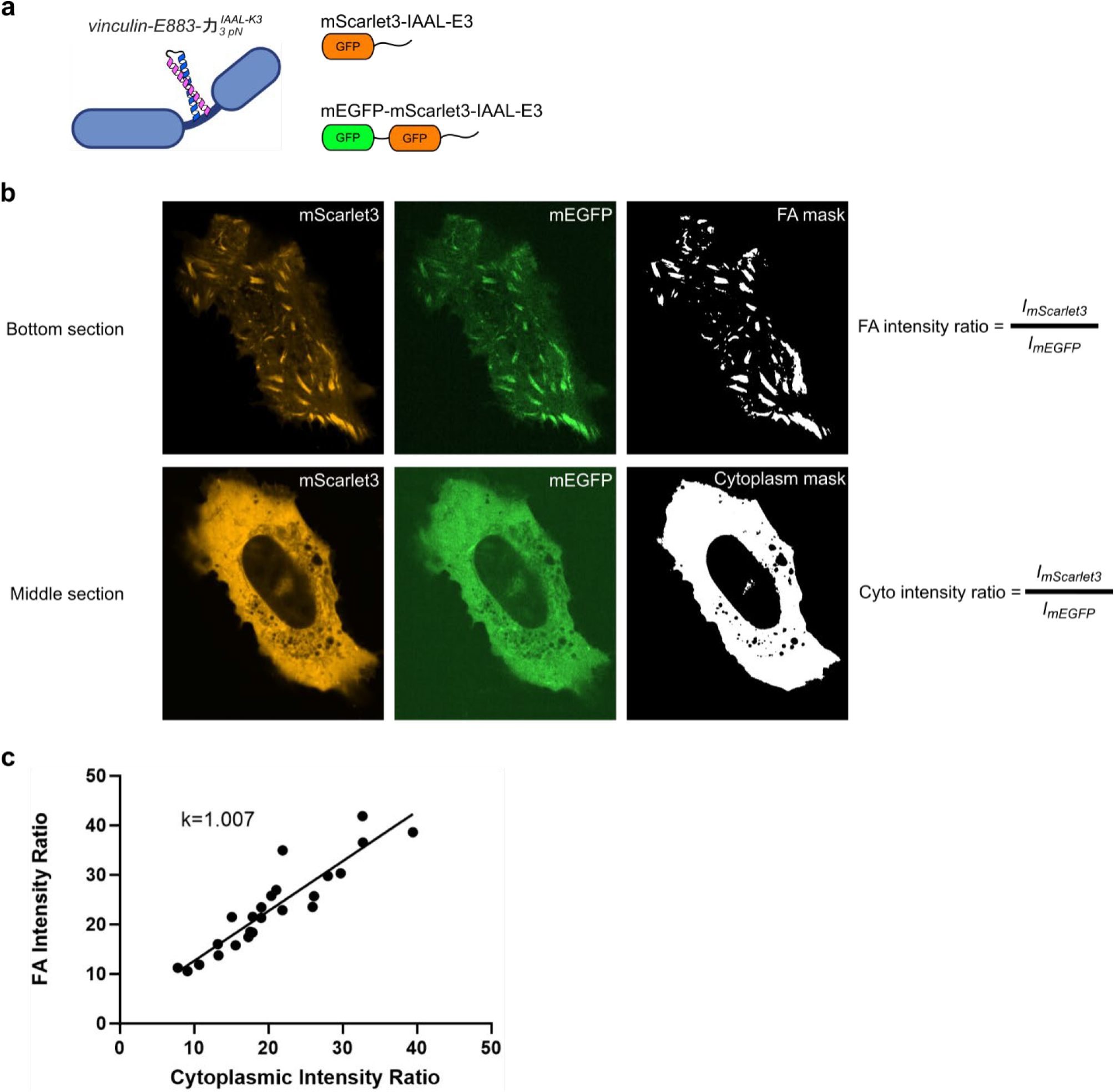
The size of the binder probe does not impede its localization to focal adhesions. **a**, U2OS cells were transfected with vinculin-E883-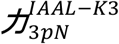 and two binders, mScarlet3-IAAL-E3 (pJB417) and mEGFP-mScarlet3-IAAL-E3 (pJB439) which is roughly twice as large. **b**, Fluorescent images from adherent U2OS cells were optically sectioned into the bottom and the middle layers. In the bottom section, images were further segmented to only record signals from focal adhesions (FA). In the middle layer, segmented signals come from the cytoplasm where mEGFP-mScarlet3-IAAL-E3 and mScarlet3-IAAL-E3 are presumably evenly distributed. We expect that if the diffusion of the probe is impeded by the other proteins in the FA, mEGFP-mScarlet3-IAAL-E3 diffusion would be more impeded than mScarlet3-IAAL-E3’s because it is two times larger, and the intensity ratio of mScarlet3 to mEGFP in the FA and in the cytoplasm would be different. **c**, Correlation of the intensity ration between FA and the cytoplasm. Each dot represents a cell. Despite the difference in variations of expression, the intensity ratios are linearly correlated, indicating no difference of FA penetration between the binders of different sizes.

**Fig. S11.**
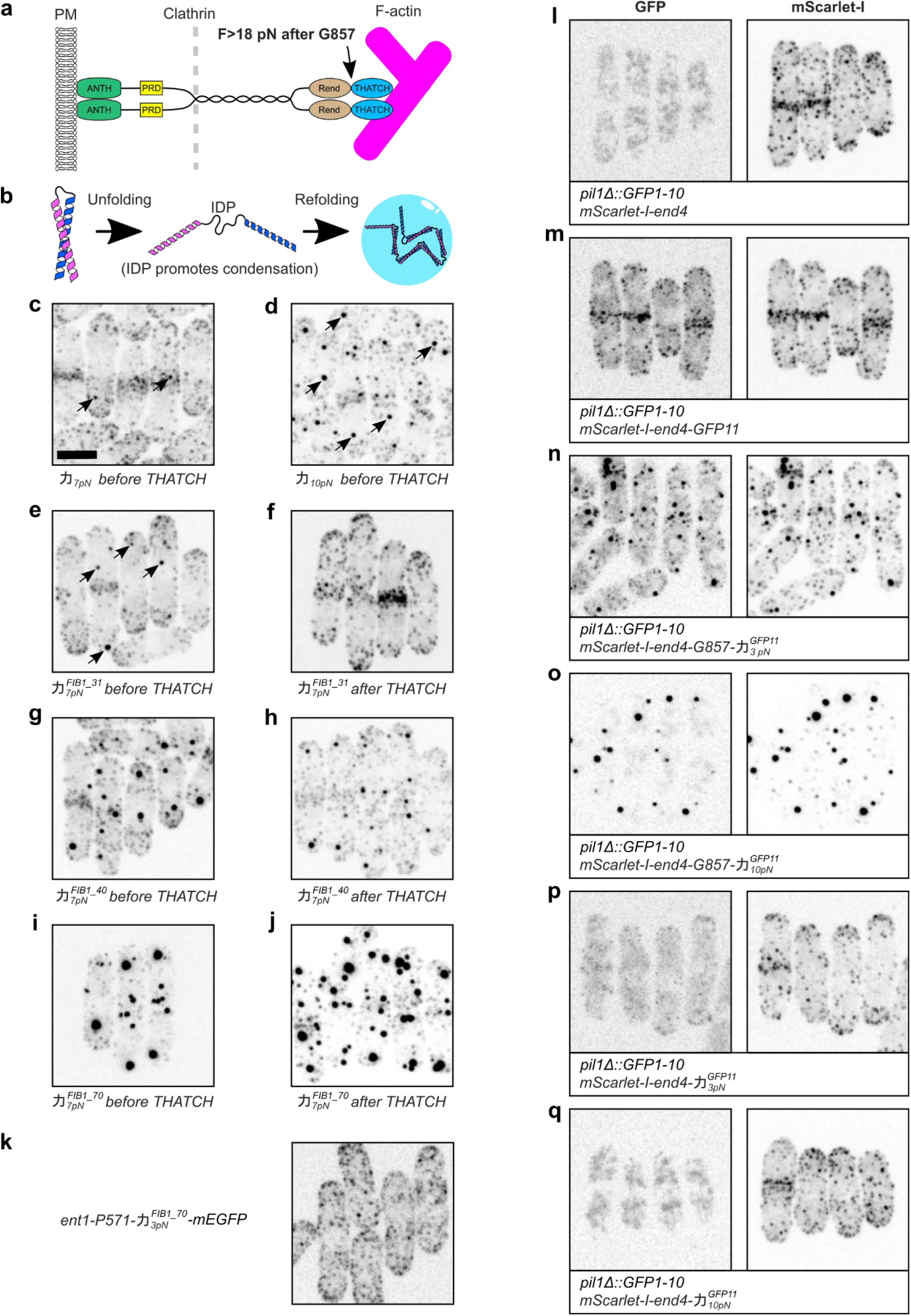
Coiled coils with IDP linkers promote the formation of protein condensates in a force-dependent manner for End4 but not Ent1. **a**, Schematic of End4, an adaptor protein that transmits force from the F-actin cytoskeleton to the membrane during clathrin-mediated endocytosis in the fission yeast. Force before THATCH is above 18 pN^27^. ANTH: AP180 N-terminal homology domain. PRD: proline rich domain. Rend: Rod in End4 domain. THATCH: talin-HIP1/R/Sla2 actin-tethering C-terminal homology domain. Not drawn to scale. **b**, Force-dependent unfolding of coiled coils exposes the IDP linker, which promotes the formation of protein condensates during the refolding of the coiled coils. **c**-**j**, End4 was fluorescently tagged, and different coiled coils were inserted into End4 either before THATCH (where F>18 pN) or after THATCH (no force control). **c**, The insertion of 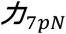 into End4 before THATCH rarely drove the formation of protein condensates (arrows) in fission yeast cells, presumably due to the relatively low affinity between the two α-helices of 力_7*pN*_. **d**, The insertion of 力 _10*pN*_ into End4 before THATCH led to the formation of large End4 condensates (arrows). **e-f**, The incorporation of the first 31 amino acids of FIB1 (FIB1_31) as the linker for 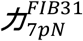 enhanced the formation of End4 condensates when inserted before THATCH (**e**) but not after THATCH (**f**). **g-h**, When a longer IDP (FIB1_40) was used as the linker of 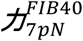, End4p condensates formed even when 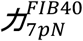 was inserted after THATCH (**h**). The effect of force can be observed by comparing the sizes of End4 condensates (larger condensates in **g** than **h**). When the IDP linker is sufficiently long (FIB1_70), the sizes of End4 condensates are comparable regardless of the site of insertion for 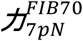. **k**,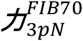 failed to promote the formation of protein condensates for Ent1 after P571, a position where the force is 5-7 pN according to the new readouts introduced in this paper. This indicates that the length of IDP required to drive protein condensation is different for different proteins, and that protein condensation is not a universal readout for force measurement. **l-q**, GFP11 can be used as a functional linker to measure force on End4. **l**, Fission yeast cells expressing GFP1-10 and mScarlet-I-end4 have fluorescent signals in the mScarlet-I channel but not the GFP channel. **m**, Starting from the strain in **l**, the insertion of GFP11 linker into End4 after G857 (flexible linker before THATCH) labeled End4p in the GFP channel. **n-o**, The insertion of either 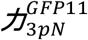 or 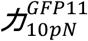 into End4 before THATCH gave dual readouts: GFP fluorescence and the formation of End4 condensates. **p-q**, The insertion of 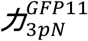 or 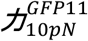 into End4 after THATCH did not lead to GFP fluorescence or End4 condensates. Images are maximum projections from confocal sections of whole cells. **l-q** are acquired and displayed using the same settings. Scale bar in **c** applies to all images, 5 μm.

## Note S1. Kinetic model of the coiled coil folding/unfolding and reporter binding

Suppose the coiled coil undergoes reversible two-state folding and unfolding transitions, and the fluorescent reporter protein only binds to the unfolded coiled coil (Fig. SN1A).

Let *p*_*f*_, *p*_*u*_, and *p*_*b*_denote the probabilities of the coiled coil being in the folded state, the free unfolded state (i.e. not bound to the reporter), and the reporter-bound state, respectively. While the force modulates the coiled coil unfolding and refolding rates (*k*_*u*_and *k*_*f*_), it is assumed that the rates of fluorescent reporter binding to and dissociating from the unfolded coiled coil (*k*_*b*_and *k*_*d*_) are approximately independent of force. Note that the reporter binding rate constant *k*_*b*_ is equal to the bimolecular association rate constant *k*_*on*_ multiplied by the reporter concentration *c* in the cell.

The state probabilities at any time *t* are related to these rate constants by the Master equations:

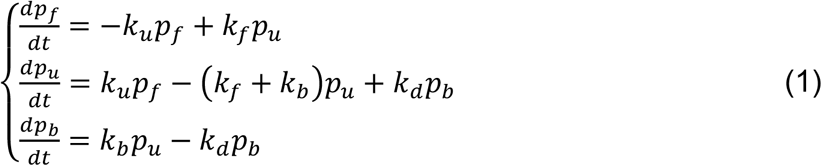

Since at any time

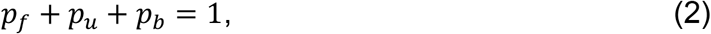

we can eliminate *p*_*f*_ in Eq. (1), yielding

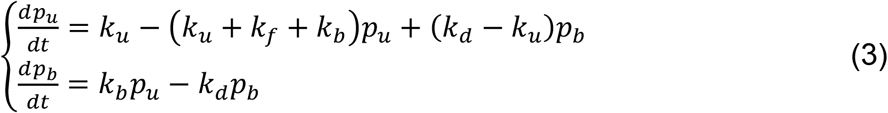

The solution of this equation set is

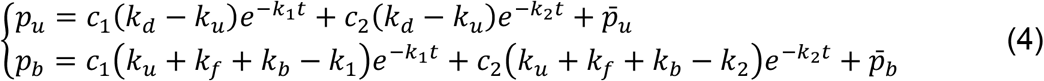

where

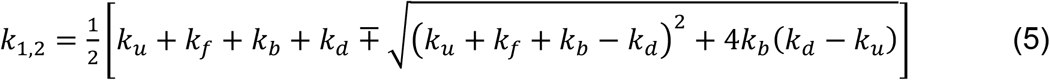

are two rate constants, and

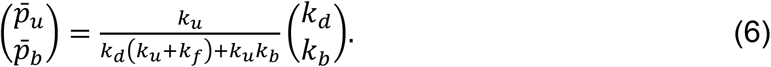

is the equilibrium solution to Eq. (3) when *t* → +∞, and *c*_2_ and *c*_2_ are two constants determined by the initial condition at *t* = 0. With the definition of the coiled coil unfolding equilibrium constant

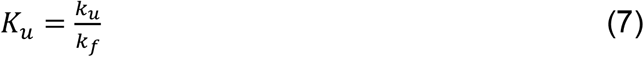

and the reporter binding equilibrium constant

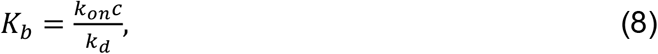

the equilibrium probabilities in Eq. (6) can be rewritten as

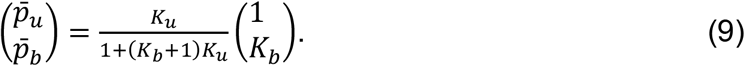

If *K*_*b*_ ≪ 1, Eq. (9) can be approximated as

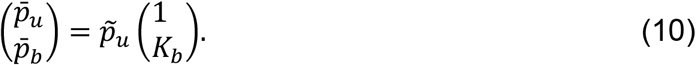

where 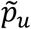 is the probability of the unfolded coiled coil in the absence of reporters, with

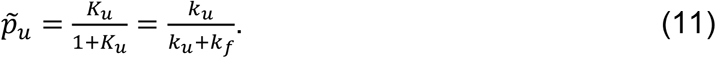

Therefore, in the case of a small equilibrium binding constant for the reporter, the probability of the reporter-bound coiled coil is proportional to the probability of unfolded coiled coil in the absence of reporter at the same force. In other words, reporter binding minimally perturbs the coiled coil unfolding.

The folding and unfolding rates of the coiled coil generally depend exponentially on force, especially around the equilibrium force (1, 2), or

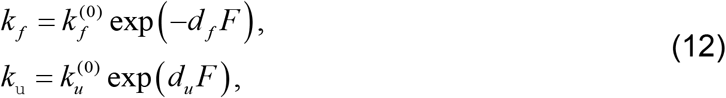

and

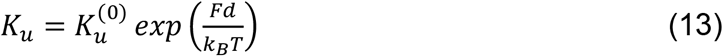

with *d* = *d*_*u*_ + *d*_*f*_, where 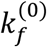 and 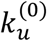 are the protein folding and unfolding rates at zero force, and 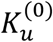 is the corresponding unfolding equilibrium constant.

### Signal to noise ratio

Suppose we image the fluorescence of force sensors in the cell using confocal fluorescence microscopy, and the sensor has a concentration or density of *ρ*_*s*_, then the fluorescence intensity of these sensors is

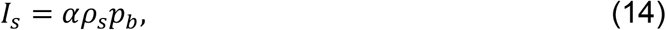

where *α*is a constant related to the properties of the fluorescence microscope and the fluorophore. Similarly, the background fluorescence from the unbound reporters is

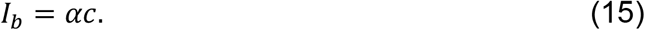

Assuming that the fluctuation of the fluorescence intensity follows a Poisson distribution, the signal signal-to-noise ratio (S/N) is

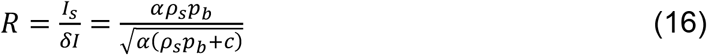

and

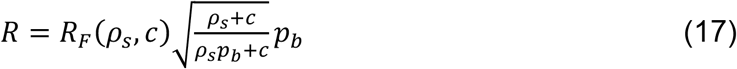

where

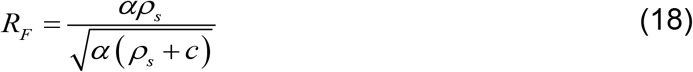

is the S/N of the microscope when detecting a fully labelled force sensor-harboring protein at density *ρ*_*s*_in the presence of free reporters with a concentration *c*in the cell.

Supposing the forces on the sensor are at equilibrium, the probability of the reporter-bound sensor *p*_*b*_in Eq. (17) can be approximated by 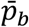 as expressed in Eq. (9), leading to 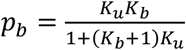 and

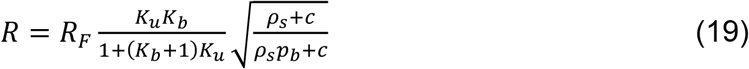

In other words, the S/N is equal to the microscope’s S/N reduced by a factor 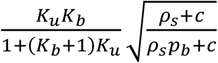.

After calculating the derivative of R with respect to c, we obtain the reporter concentration *c*_*Rmax*_ that maximizes the S/N:

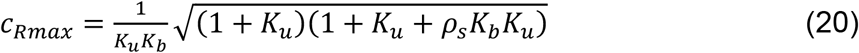

### Sudden force drops and force jumps

Now let’s examine the temporal response of the reporter to a force jump or drop in two extreme cases. In the first case, the sensor is subject to a force drop to zero. Because in the absence of force the folding rate of the coiled coil is generally much greater than other rates, or *k*_*f*_ ≫ *k*_*u*_, *k*_*b*_, *k*_*d*_, we can approximate Eq. (5) as

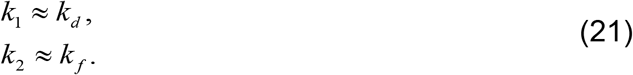

Therefore, for a fast-vanishing force, the response of the force sensor is limited by the dissociation rate of the reporter.

In the second case, the sensor is subject to a large force jump, such that at the high force, *k*_*u*_ ≫ *k*_*f*_, *k*_*b*_, *k*_*d*_. In this case,

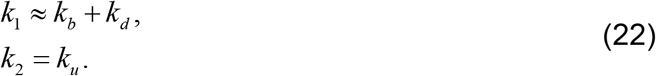

Therefore, for a fast and significant force jump, the response of the force sensor is limited by the sum of the reporter binding and dissociation rates.

### Simulations

To illustrate these results, we chose parameters to represent our force sensor with an equilibrium force of *F*_1*/*2_ = 10 pN, an extension change *d* = 10nm, and a folding rate 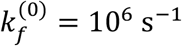 at zero force. Correspondingly, the unfolding energy of the coiled coil *V* = *F*_1/2_*d* = 24.4 k_B_T. The unfolding rate was calculated as 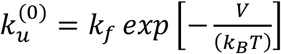 The bimolecular binding rate constant was chosen as 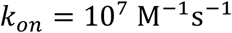, approaching the diffusion-limited bimolecular association rate. The live reporter readout IAAL-K3/ mEGFP-IAAL-E3 has a high binding affinity of 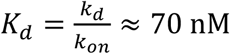 measured *in vitro*. However, the actual *in vivo* dissociation constant may differ from this value, due to molecular crowding effects. Therefore, we considered a range of reporter binding affinities in the 70-200 nM range. The reporter dissociation rate was calculated as *k*_*d*_ = *K*_*d*_*k*_*on*_. Finally, the reporter concentration in the cell used in our assay was estimated to be *c* = 1 µM. If not otherwise specified, these parameters served as default parameters for our calculations.

We first calculated the equilibrium probabilities of the three sensor states as a function of the force applied to the sensor, with a reporter binding affinity 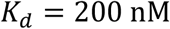 or binding equilibrium constant 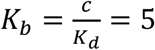, using Eqs. (9) and (13). As the force increases, the coiled coil rapidly unfolds at 8-10.5 pN and is concurrently bound by the reporter (Fig. SN1B, top panel). While the probability of the folded coiled coil becomes negligible above 11.5 pN, the probability of the reporter-bound state reaches a plateau of 0.83. Correspondingly, the probability of the unfolded coiled coil state free of reporter binding plateaus at 0.17. The sigmoidal profile of the reporter-binding probability is close to that of the unfolding probability of the coiled coil only, but slightly shifts toward lower force by 0.7 pN, as judged by the force values at half the equilibrium probability (Fig. SN1B, middle panel). A decrease in the reporter concentration to 0.1 µM with *K*_*b*_ = 1 reduces the force shift to 0.3 pN (data not shown). In contrast, increasing the reporter binding affinity to *K*_*d*_ = 70 nM or *K*_*b*_ = 14.3 lowers the binding probability profile by 1.1 pN. Therefore, the reporter binding slightly shifts the protein unfolding equilibrium to lower force by a few percents in a predictable *K*_*b*_ -dependent manner.

Next, we examined the ability of an imaging system to detect an open force sensor. The S/N principally depends on the sensitivity of the fluorescence microscope used to image the force sensors, and is typically in the range of 10-100. For the rest of the calculations, we chose a microscope S/N such that *R*_*F*_ = 10 using the default parameters. According to Eq. (20), the S/N of our microscope is reduced by a factor that depends on the biochemical and biophysical properties of the sensor and the reporter. Using a high affinity reporter (Kd<200 nM) only marginally reduces the S/N and using a lower affinity reporter reduces the S/N by a factor 2 or more if the reporter concentration is also low. Our calculations and simulations also demonstrate that one can maximize the S/N by choosing an optimum reporter concentration (e.g. 490 nM with the default parameters), and the S/N decreases slowly with increasing reporter concentrations past this optimum. Ultimately the overall S/N of force detection depends on the microscope’s S/N and using a more sensitive fluorescence microscope can compensate for the effect of a low affinity reporter system.

Next, we determined the temporal resolution of our force sensor to detect fast force changes. The time-dependent probability of the reporter-bound coiled coil was calculated as the coiled coil is subject to a force jump from zero force (t<0) to a specified force (t>0, Fig. SN1C, top panel). The coiled coil is held at the force for 5 seconds and then completely relaxed. Upon the force jump, the probability increases with a rate that increases with the force (Fig. SN1C, middle panel). At 8 pN force, the probability for the coiled coil to be unfolded grows slowly, leading to slow reporter binding and low binding probability. At forces equal to or greater than the 10 pN equilibrium force of the coiled coil, the reporter binds much more quickly, with the greatest rate at 14 pN. A decrease in *K*_*d*_ from 200 nM to 70 nM does not significantly change the reporter binding rates. At 14 pN, the reporter binding rates at both reporter binding affinities become limited by the sum of the reporter binding and dissociation rates *k*_1_ = 12 *s*^−1^, as predicted by Eq. (23). A higher affinity slightly promotes the final reporter binding at all force at t=5 s, as expected. Finally, the fast protein folding significantly contributes to the fast force response. A 10-fold decrease in the folding rate *k*_*f*_ reduces the rate of the sensor response to the 14 pN force jump to *k*_1_ = 4 *s*^−1^ (data not shown).

The force sensor responds to a drop in force from different initial levels with an identical decay rate, corresponding to the reporter dissociation rate *k*_*d*_ = 2 *s*^−1^, as predicted by Eq. (22) (Fig. SN1C, middle panel). In these calculations, we chose the equilibrium condition at high force at t=5 s as the initial state for the force drop. Reducing the reporter’s dissociation constant from 200 nM to 70 nM decreases the decay rate to 0.7 s^-1^ (Fig. SN1C, bottom panel). Therefore, given a constant reporter binding rate *k*_*b*_, a weaker reporter binding affinity leads to a faster sensor response to force decreases.

Rapid, force-dependent coiled coil folding/unfolding transition and fast reporter binding are essential for capturing rapid force increases, while a high reporter dissociation rate is necessary for detecting force decreases. Coiled coils are particularly well suited for force sensing due to their rapid folding and force-sensitive transition rates at the equilibrium force. Given that the binding rate is constrained by the diffusion-limited bimolecular binding (∼10^8^ M^-1^s^-1^) and the need to limit intracellular reporter concentration (to avoid increased fluorescence background), lower reporter affinity is preferable for detecting fast force dynamics.

From a practical standpoint, our calculations suggest that when the force applied is significantly different from the sensor’s nominal threshold, the force sensors will report the correct amount of force within a few percents of the sensor’s threshold and on a subsecond timescale, for realistic rate constants of folding, unfolding and reporter binding and unbinding. Therefore, it is suggested to collect data with several sensors with different force thresholds, and, if force is expected to be constant, avoid over-interpreting the kinetics data obtained with sensors with nominal thresholds very close to this applied force.

**Fig. SN1.**
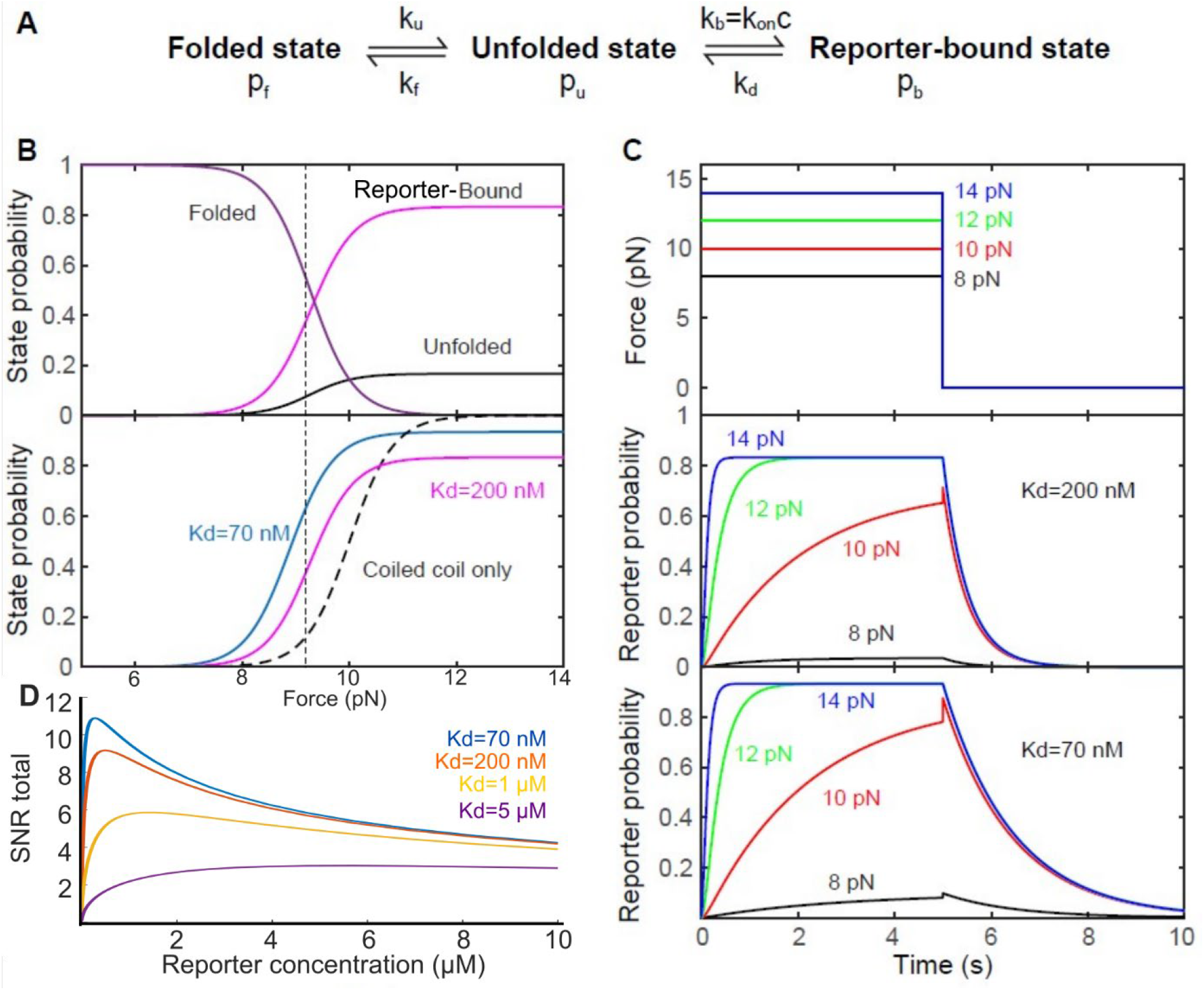
Theoretical model illustrating the resolution of force sensors based on force-induced protein unfolding and refolding. (A) Protein conformational states involved in force sensing, with association transitions rates and state probabilities. (B) Force-dependent state probability. (C) Time-dependent force (top panel) and probabilities of reporter-bound coiled coils (middle and bottom panels). (D) Microscope’s SNR when using force sensors with live reporters of different affinities.

## Notes

### Summary of Updates

- Measurements on vinculin were extended, including FRAP, simulatneous use of 2 sensors, cross-validation between readouts - Force spectroscopy data of coiled-coil opening at different pulling speed, or with different linkers were added - Figures were reorganized - A supplemental note with a model for the kinetics and detection SNR of the sensors was added

## References

1. Wang, X. & Ha, T. Defining single molecular forces required to activate integrin and notch signaling. Science 340, 991–994 (2013).

2. Blakely, B. L. et al. A DNA-based molecular probe for optically reporting cellular traction forces. Nat Methods 11, 1229–1232 (2014).

3. Zhang, Y., Ge, C., Zhu, C. & Salaita, K. DNA-based digital tension probes reveal integrin forces during early cell adhesion. Nat Commun 5, 5167 (2014).

4. Grashoff, C. et al. Measuring mechanical tension across vinculin reveals regulation of focal adhesion dynamics. Nature 466, 263 (2010).

5. Suzuki, A. et al. How the kinetochore couples microtubule force and centromere stretch to move chromosomes. Nat Cell Biol 18, 382–392 (2016).

6. Amiri, S. et al. Intracellular tension sensor reveals mechanical anisotropy of the actin cytoskeleton. Nat Commun 14, 8011 (2023).

7. Polacheck, W. J. & Chen, C. S. Measuring cell-generated forces: a guide to the available tools. Nat Methods 13, 415–423 (2016).

8. Jurchenko, C. & Salaita, K. S. Lighting Up the Force: Investigating Mechanisms of Mechanotransduction Using Fluorescent Tension Probes. Mol Cell Biol 35, 2570–2582 (2015).

9. LaCroix, A. S., Rothenberg, K. E., Berginski, M. E., Urs, A. N. & Hoffman, B. D. Chapter 10 - Construction, imaging, and analysis of FRET-based tension sensors in living cells. in Methods in Cell Biology (ed. Paluch, E. K.) vol. 125 161–186 (Academic Press, 2015).

10. Ham, T. R., Collins, K. L. & Hoffman, B. D. Molecular Tension Sensors: Moving Beyond Force. Curr Opin Biomed Eng 12, 83–94 (2019).

11. LaCroix, A. S., Lynch, A. D., Berginski, M. E. & Hoffman, B. D. Tunable molecular tension sensors reveal extension-based control of vinculin loading. eLife 7, e33927 (2018).

12. Ringer, P. et al. Multiplexing molecular tension sensors reveals piconewton force gradient across talin-1. Nature Methods 14, 1090–1096 (2017).

13. Lemke, S. B., Weidemann, T., Cost, A.-L., Grashoff, C. & Schnorrer, F. A small proportion of Talin molecules transmit forces at developing muscle attachments in vivo. PLoS Biol 17, e3000057 (2019).

14. Jo, M. H. et al. Determination of single-molecule loading rate during mechanotransduction in cell adhesion. Science 383, 1374–1379 (2024).

15. Hu, Y. et al. DNA-based ForceChrono probes for deciphering single-molecule force dynamics in living cells. Cell 0, (2024).

16. Abella, M., Andruck, L., Malengo, G. & Skruzny, M. Actin-generated force applied during endocytosis measured by Sla2-based FRET tension sensors. Developmental Cell 56, 2419–2426.e4 (2021).

17. Liu, J. et al. Tension Gauge Tethers as Tension Threshold and Duration Sensors. ACS Sens. 8, 704–711 (2023).

18. Mosayebi, M., Louis, A. A., Doye, J. P. K. & Ouldridge, T. E. Force-Induced Rupture of a DNA Duplex: From Fundamentals to Force Sensors. ACS Nano 9, 11993–12003 (2015).

19. Cocco, S., Monasson, R. & Marko, J. F. Force and kinetic barriers to unzipping of the DNA double helix. Proceedings of the National Academy of Sciences 98, 8608–8613 (2001).

20. Sun, X., Hao, P. & Wu, N. DNA-Based Mechanical Sensors for Cell Applications. Chemistry 5, 1546–1559 (2023).

21. Brockman, J. M. et al. Live-cell super-resolved PAINT imaging of piconewton cellular traction forces. Nat Methods 17, 1018–1024 (2020).

22. Liu, Y. et al. DNA-based nanoparticle tension sensors reveal that T-cell receptors transmit defined pN forces to their antigens for enhanced fidelity. Proceedings of the National Academy of Sciences 113, 5610–5615 (2016).

23. Aird, E. J., Tompkins, K. J., Ramirez, M. P. & Gordon, W. R. Enhanced Molecular Tension Sensor Based on Bioluminescence Resonance Energy Transfer (BRET). ACS Sens. 5, 34–39 (2020).

24. Cost, A.-L., Ringer, P., Anna, C.-G. & Grashoff, C. How to measure molecular forces in cells: a guide to evaluating genetically-encoded FRET-based tension sensors. 8, 96–105 (2015).

25. Eder, D., Basler, K. & reports, A.-C. Challenging FRET-based E-Cadherin force measurements in Drosophila. Scientific reports (2017).

26. As, L., Ke, R., Me, B., An, U. & Bd, H. Construction, imaging, and analysis of FRET-based tension sensors in living cells. Methods in cell biology 125, (2015).

27. Ren, Y. et al. Force redistribution in clathrin-mediated endocytosis revealed by coiled-coil force sensors. Sci Adv 9, eadi1535 (2023).

28. Ren, Y., Yang, J., Jin, H., Zhang, Y. & Berro, J. Force Redistribution in Clathrin-Mediated Endocytosis Revealed by Phase-Separating Force Sensors. 2021.06.29.450294 https://www.biorxiv.org/content/10.1101/2021.06.29.450294v3 (2021) xdoi:10.1101/2021.06.29.450294.

29. Berro, J., Ren, Y. & Zhang, Y. Coiled-coil peptides for force-dependent applications. (2024).

30. Austen, K. et al. Extracellular rigidity sensing by talin isoform-specific mechanical linkages. Nature Cell Biology 17, 1597–1606 (2015).

31. Zhong, B. L., Vachharajani, V. T. & Dunn, A. R. STReTCh: A Strategy for Facile Detection of Mechanical Forces across Proteins in Cells. http://biorxiv.org/lookup/doi/10.1101/2021.12.31.474658 (2022) xdoi:10.1101/2021.12.31.474658.

32. Zhong, B. L. et al. Split Luciferase Molecular Tension Sensors for Bioluminescent Readout of Mechanical Forces in Biological Systems. ACS Sens. 9, 3489–3495 (2024).

33. McClain, D. L., Woods, H. L. & Oakley, M. G. Design and Characterization of a Heterodimeric Coiled Coil that Forms Exclusively with an Antiparallel Relative Helix Orientation. J. Am. Chem. Soc. 123, 3151–3152 (2001).

34. Park, W. M. Coiled-Coils: The Molecular Zippers that Self-Assemble Protein Nanostructures. Int J Mol Sci 21, 3584 (2020).

35. Truebestein, L. & Leonard, T. A. Coiled-coils: The long and short of it. Bioessays 38, 903–916 (2016).

36. Su, J. Y., Hodges, R. S. & Kay, C. M. Effect of Chain Length on the Formation and Stability of Synthetic.alpha.-Helical Coiled Coils. Biochemistry 33, 15501–15510 (1994).

37. Litowski, J. r. & Hodges, R. s. Designing heterodimeric two-stranded α-helical coiled-coils: the effect of chain length on protein folding, stability and specificity. The Journal of Peptide Research 58, 477–492 (2001).

38. Carrion-Vazquez, M. et al. Mechanical and chemical unfolding of a single protein: A comparison. Proc. Natl. Acad. Sci. U.S.A. 96, 3694–3699 (1999).

39. Evans, E. Probing the Relation Between Force—Lifetime—and Chemistry in Single Molecular Bonds. Annual Review of Biophysics 30, 105–128 (2001).

40. Biewenga, L., Rosier, B. J. H. M. & Merkx, M. Engineering with NanoLuc: a playground for the development of bioluminescent protein switches and sensors. Biochemical Society Transactions 48, 2643–2655 (2020).

41. Schwinn, M. K. et al. CRISPR-Mediated Tagging of Endogenous Proteins with a Luminescent Peptide. ACS Chem. Biol. 13, 467–474 (2018).

42. Kamiyama, D. et al. Versatile protein tagging in cells with split fluorescent protein. Nature Communications 7, 11046 (2016).

43. Romei, M. G. & Boxer, S. G. Split Green Fluorescent Proteins: Scope, Limitations, and Outlook. Annu Rev Biophys 48, 19–44 (2019).

44. Raran-Kurussi, S., Cherry, S., Zhang, D. & Waugh, D. S. Removal of Affinity Tags with TEV Protease. Methods Mol Biol 1586, 221–230 (2017).

45. Lindhout, D. A., Litowski, J. R., Mercier, P., Hodges, R. S. & Sykes, B. D. NMR solution structure of a highly stable de novo heterodimeric coiled-coil. Biopolymers 75, 367–375 (2004).

46. Liu, B. et al. Biosensors based on peptide exposure show single molecule conformations in live cells. Cell 184, 5670–5685.e23 (2021).

47. Shekhar, S. et al. Formin and capping protein together embrace the actin filament in a ménage à trois. Nature Communications 6, 8730 (2015).

48. Coffman, V. C., Sees, J. A., Kovar, D. R. & Wu, J.-Q. The formins Cdc12 and For3 cooperate during contractile ring assembly in cytokinesis. J Cell Biol 203, 101–114 (2013).

49. Jégou, A., Carlier, M.-F. & Romet-Lemonne, G. Formin mDia1 senses and generates mechanical forces on actin filaments. Nat Commun 4, 1883 (2013).

50. Courtemanche, N., Lee, J. Y., Pollard, T. D. & Greene, E. C. Tension modulates actin filament polymerization mediated by formin and profilin. Proc Natl Acad Sci U S A 110, 9752–9757 (2013).

51. Yu, M. et al. Effects of Mechanical Stimuli on Profilin- and Formin-Mediated Actin Polymerization. Nano Lett 18, 5239–5247 (2018).

52. Yu, M. et al. mDia1 senses both force and torque during F-actin filament polymerization. Nat Commun 8, 1650 (2017).

53. Saito, T., Ren, Y. & Berro, J. Mechanisms of force transmission on cytokinesis Formin Cdc12 in fission yeast revealed by coiled-coil force sensors. 2025.05.14.653946 Preprint at 10.1101/2025.05.14.653946 (2025).

54. Aguilar, R. C., Watson, H. A. & Wendland, B. The Yeast Epsin Ent1 Is Recruited to Membranes through Multiple Independent Interactions. J. Biol. Chem. 278, 10737–10743 (2003).

55. Skruzny, M. et al. Molecular basis for coupling the plasma membrane to the actin cytoskeleton during clathrin-mediated endocytosis. Proc Natl Acad Sci U S A 109, E2533–E2542 (2012).

56. Picco, A., Mund, M., Ries, J., Nédélec, F. & Kaksonen, M. Visualizing the functional architecture of the endocytic machinery. eLife 4, e04535.

57. Berro, J. & Pollard, T. D. Local and global analysis of endocytic patch dynamics in fission yeast using a new ‘temporal superresolution’ realignment method. Mol Biol Cell 25, 3501–3514 (2014).

58. Lemière, J., Ren, Y. & Berro, J. Rapid adaptation of endocytosis, exocytosis, and eisosomes after an acute increase in membrane tension in yeast cells. eLife 10, e62084 (2021).

59. Boulant, S., Kural, C., Zeeh, J.-C., Ubelmann, F. & Kirchhausen, T. Actin dynamics counteract membrane tension during clathrin-mediated endocytosis. Nat Cell Biol 13, 1124–1131 (2011).

60. Lacy, M. M., Ma, R., Ravindra, N. G. & Berro, J. Molecular mechanisms of force production in clathrin-mediated endocytosis. FEBS Lett. 592, 3586–3605 (2018).

61. Skruzny, M., Pohl, E., Gnoth, S., Malengo, G. & Sourjik, V. The protein architecture of the endocytic coat analyzed by FRET microscopy. Molecular Systems Biology 16, e9009 (2020).

62. Unsain, N., Stefani, F. D. & Cáceres, A. The Actin/Spectrin Membrane-Associated Periodic Skeleton in Neurons. Front. Synaptic Neurosci. 10, (2018).

63. Costa, A. R. et al. The membrane periodic skeleton is an actomyosin network that regulates axonal diameter and conduction. eLife 9, e55471 (2020).

64. Glomb, O. et al. A kinesin-1 adaptor complex controls bimodal slow axonal transport of spectrin in Caenorhabditis elegans. Dev Cell 58, 1847–1863.e12 (2023).

65. Lundquist, E. A., Herman, R. K., Shaw, J. E. & Bargmann, C. I. UNC-115, a Conserved Protein with Predicted LIM and Actin-Binding Domains, Mediates Axon Guidance in C. elegans. Neuron 21, 385–392 (1998).

66. Zhou, R. et al. Proteomic and functional analyses of the periodic membrane skeleton in neurons. Nat Commun 13, 3196 (2022).

67. Krieg, M., Dunn, A. R. & Goodman, M. B. Mechanical Control of the Sense of Touch by β Spectrin. Nat Cell Biol 16, 224–233 (2014).

68. Crisp, M. et al. Coupling of the nucleus and cytoplasm: role of the LINC complex. J Cell Biol 172, 41–53 (2006).

69. Gottardi, C. J. & Luxton, G. W. G. Nesprin-2G tension fine-tunes Wnt/β-catenin signaling. J Cell Biol 219, e202009042 (2020).

70. Srivastava, L. K. & Ehrlicher, A. J. Sensing the squeeze: nuclear mechanotransduction in health and disease. Nucleus 15, 2374854 (2024).

71. Arsenovic, P. T. et al. Nesprin-2G, a Component of the Nuclear LINC Complex, Is Subject to Myosin-Dependent Tension. Biophys J 110, 34–43 (2016).

72. Shams, H., Luxton, G. & Mofrad, M. R. K. Mechanical Response of Nesprin-2G to Cytoskeletal Forces Regulates Linc Complex-Dependent Mechanotransduction. Biophysical Journal 112, 337a (2017).

73. Arsenovic, P. T. et al. Nesprin-2G, a Component of the Nuclear LINC Complex, Is Subject to Myosin-Dependent Tension. Biophys J 110, 34–43 (2016).

74. Carley, E. et al. The LINC complex transmits integrin-dependent tension to the nuclear lamina and represses epidermal differentiation. eLife 10, e58541 (2021).

75. Rothenberg, K. E., Scott, D. W., Christoforou, N. & Hoffman, B. D. Vinculin Force-Sensitive Dynamics at Focal Adhesions Enable Effective Directed Cell Migration. Biophys J 114, 1680–1694 (2018).

76. Liu, X. et al. The mechanical response of vinculin. 2023.05.25.542235 Preprint at 10.1101/2023.05.25.542235 (2023).

77. Huang, D. L., Bax, N. A., Buckley, C. D., Weis, W. I. & Dunn, A. R. Vinculin forms a directionally asymmetric catch bond with F-actin. Science 357, 703–706 (2017).

78. Kumar, A. et al. Talin tension sensor reveals novel features of focal adhesion force transmission and mechanosensitivity. Journal of Cell Biology 213, 371–383 (2016).

79. Wang, Y. et al. Force-Dependent Interactions between Talin and Full-Length Vinculin. J. Am. Chem. Soc. 143, 14726–14737 (2021).

80. Yao, M. et al. Mechanical activation of vinculin binding to talin locks talin in an unfolded conformation. Sci Rep 4, 4610 (2014).

81. Ren, Y., Yang, J., Fujita, B., Zhang, Y. & Berro, J. Cross-regulations of two connected domains form a mechanical circuit for steady force transmission during clathrin-mediated endocytosis. Cell Rep 43, 114725 (2024).

82. Wang, Y., Yan, J. & Goult, B. T. Force-Dependent Binding Constants. Biochemistry (2019) doi:10.1021/acs.biochem.9b00453.

83. Naughton, B. S. et al. Nodal Modulator (NOMO) is a force-bearing transmembrane protein required for muscle differentiation. 2024.09.06.611727 Preprint at 10.1101/2024.09.06.611727 (2024).

84. Berro, J., Ren, Y. & Zhang, Y. Coiled-coil peptides for force-dependent applications. (2024).

85. Gao, Y. et al. Single reconstituted neuronal SNARE complexes zipper in three distinct stages. Science 337, 1340–1343 (2012).

86. Cecconi, C., Shank, E. A., Dahlquist, F. W., Marqusee, S. & Bustamante, C. Protein-DNA chimeras for single molecule mechanical folding studies with the optical tweezers. Eur Biophys J 37, 729–738 (2008).

87. Yang, J., Jin, H., Liu, Y., Guo, Y. & Zhang, Y. A dynamic template complex mediates Munc18-chaperoned SNARE assembly. Proceedings of the National Academy of Sciences 119, e2215124119 (2022).

88. Gao, Y., Sirinakis, G. & Zhang, Y. Highly Anisotropic Stability and Folding Kinetics of a Single Coiled Coil Protein under Mechanical Tension. J. Am. Chem. Soc. 133, 12749–12757 (2011).

89. Jiao, J., Rebane, A. A., Ma, L. & Zhang, Y. Single-Molecule Protein Folding Experiments Using High-Precision Optical Tweezers. Methods Mol Biol 1486, 357–390 (2017).

90. Jumper, J. et al. Highly accurate protein structure prediction with AlphaFold. Nature 596, 583–589 (2021).

91. Goktas, M. et al. Molecular mechanics of coiled coils loaded in the shear geometry †Electronic supplementary information (ESI) available: CD measurements for the determination of secondary structure and the thermal stability of the coiled coils as well as additional results of the SMFS experiments and the SMD simulations are included in the supporting information. See DOI: 10.1039/c8sc01037d. Chem Sci 9, 4610–4621 (2018).

92. Fernandez, R. & Berro, J. Use of a fluoride channel as a new selection marker for fission yeast plasmids and application to fast genome editing with CRISPR/Cas9. Yeast 33, 549–557 (2016).

93. Fernandez, R. & Berro, J. CRISPR-Cas9 editing efficiency in fission yeast is not limited by homology search and is improved by combining gap-repair with fluoride selection. 2024.03.01.582946 Preprint at 10.1101/2024.03.01.582946 (2024).

94. Wood, V. et al. PomBase: a comprehensive online resource for fission yeast. Nucleic Acids Research 40, D695–D699 (2012).

95. Schindelin, J. et al. Fiji: an open-source platform for biological-image analysis. Nat Methods 9, 676–682 (2012).

96. Lacy, M. M., Baddeley, D. & Berro, J. Single-molecule turnover dynamics of actin and membrane coat proteins in clathrin-mediated endocytosis. eLife 8, e52355 (2019).

97. Mousavi, S. I., Lacy, M. M., Li, X. & Berro, J. Fast Actin Disassembly and Fimbrin Mechanosensitivity Support Rapid Turnover in a Model of Clathrin-Mediated Endocytosis. Cytoskeleton (Hoboken) (2025) doi:10.1002/cm.22002.

98. Schneider, C. A., Rasband, W. S. & Eliceiri, K. W. NIH Image to ImageJ: 25 years of image analysis. Nat Methods 9, 671–675 (2012).

99. Glomb, O., Lyu, M. & Yogev, S. Optimizing Visualization of Axonal Transport of Endogenous Cargo by Fluorescence Microscopy in Living Caenorhabditis elegans. J Vis Exp (2024) doi:10.3791/66236.

100. Preibisch, S., Saalfeld, S. & Tomancak, P. Globally optimal stitching of tiled 3D microscopic image acquisitions. Bioinformatics 25, 1463–1465 (2009).

101. Östlund, C. et al. Dynamics and molecular interactions of linker of nucleoskeleton and cytoskeleton (LINC) complex proteins. Journal of Cell Science 122, 4099–4108 (2009).

102. Tsai, P.-L. et al. Dynamic quality control machinery that operates across compartmental borders mediates the degradation of mammalian nuclear membrane proteins. Cell Reports 41, 111675 (2022).

103. Abramson, J. et al. Accurate structure prediction of biomolecular interactions with AlphaFold 3. Nature 630, 493–500 (2024).

